# Odor representations from the two nostrils are temporally segregated in human piriform cortex

**DOI:** 10.1101/2023.02.14.528521

**Authors:** Gülce Nazlı Dikecligil, Andrew I. Yang, Nisha Sanghani, Timothy Lucas, H. Isaac Chen, Kathryn A. Davis, Jay A. Gottfried

**Affiliations:** Department of Neurology, Perelman School of Medicine, University of Pennsylvania, Philadelphia, PA, 19104, USA; Department of Neurosurgery, Barrow Neurological Institute, Phoenix, AZ, 85013, USA; Department of Neurosurgery and Biomedical Engineering, Ohio State University, Columbus, OH, 43210, USA; Department of Neurosurgery, Perelman School of Medicine, University of Pennsylvania, Philadelphia, PA, 19104, USA; Department of Psychology, School of Arts and Sciences, University of Pennsylvania, Philadelphia, PA, 19104, USA

## Abstract

The human olfactory system has two discrete channels of sensory input, arising from olfactory epithelia housed in the left and right nostrils. Here, we asked whether primary olfactory cortex (piriform cortex, PC) encodes odor information arising from the two nostrils as integrated or distinct stimuli. We recorded intracranial EEG signals directly from PC while human subjects participated in an odor identification task where odors were delivered to the left, right, or both nostrils. We analyzed the time-course of odor-identity coding using machine learning approaches, and found that uni-nostril odor inputs to the ipsilateral nostril are encoded ∼480 ms faster than odor inputs to the contralateral nostril on average. During naturalistic bi-nostril odor sampling, odor information emerged in two temporally segregated epochs with the first epoch corresponding to the ipsilateral and the second epoch corresponding to the contralateral odor representations. These findings reveal that PC maintains distinct representations of odor input from each nostril through temporal segregation, highlighting an olfactory coding scheme at the cortical level that can parse odor information across nostrils within the course of a single inhalation.

## INTRODUCTION

The human olfactory system has two discrete channels of sensory input within the nose. Olfactory epithelia in the left and right nostrils, isolated from one another by the nasal septum, each receive a snapshot of the sensory world upon inhalation. However, as odor stimuli in the natural environment can change rapidly over time and space^1–4^, the olfactory composition of these two snapshots may vary considerably on any given sniff^5^. As such, while olfactory information arising from the two nostrils ultimately converges onto a unified odor percept, studies across animals and humans have shown that inter-nostril differences can also inform and shape behavior^6–9^. For instance, differences in odor concentration across the two nostrils help rodents localize an odor source^6^ and can bias motion perception in humans towards the nostril with the higher concentration odorant^9^. Such findings strongly imply that the olfactory system is equipped to both integrate and segregate odor information arising from each side of the nasal cavity. However, despite extensive work on odor responses in the olfactory system^10–15^, relatively little is known about how information from the two nostrils is integrated and differentiated in the human olfactory system^16^. Specifically, if PC favors integration or segregation of odor information across the two nostrils remains poorly understood.

The question of sensory integration across two discrete channels is particularly interesting in the context of olfaction, as the ascending olfactory pathway from the periphery to PC is primarily organized ipsilaterally: sensory neurons in each epithelium synapse onto glomeruli in the ipsilateral OB, which then project to ipsilateral PC^17^. Despite these prominent ipsilateral projections, interhemispheric communication at different stages of the olfactory pathway is thought to underlie sensory integration across nostrils^16,18–23^. Recent work outlining the interhemispheric connections in the rodent olfactory system indicates that odor information arising from the ipsilateral nostril reaches PC in just two synapses, whereas odor information arising from the contralateral nostril takes a more circuitous route^16^, raising the intriguing possibility that odor information from the two nostrils may not be encoded in PC simultaneously.

Single neuron recordings in rodents^21,22^ and neuroimaging studies in humans^24,25^ have shown that PC can respond to odor inputs arising from either nostril, although the time course of these responses and its impact on sensory convergence across nostrils have not been investigated. Additionally, whether integration or segregation across nostrils underlies odor coding in the human PC is not known. Lastly, in contrast to rodents, relatively little is known about the neural circuitry of the human olfactory system and the pathways that support interhemispheric communication of olfactory information. One possibility is that odor information arising from each epithelium is integrated early in the processing stream, at the level of the OB (as proposed by recent work in rodents^20^), such that odor information arriving at PC has already been integrated across nostrils. Alternatively, inputs from each nostril might be represented as independent stimuli with distinct spatial and/or temporal response profiles, as a consequence of the putative differences between the ipsilateral and contralateral pathways^16^.

In this study we set out to investigate if human PC maintains distinct and separable representations of odor information arising from each nostril. To this end, we recorded intracranial EEG (iEEG) signals from the PC of epilepsy patients undergoing invasive monitoring, enabling us to characterize odor responses with high spatial and temporal resolution. Subjects participated in an odor identification task where odors were delivered either to the left, right, or bilateral nostrils via a computer-controlled olfactometer and were asked to report which odor they smelled and which nostril the odor stimuli arose from. Conducting the experiment with human subjects enabled us to combine two perceptual questions within one paradigm and assess odor percepts without extensive training, an experimental advantage that would not be feasible in animal models. This study design allowed us to investigate the temporal evolution of odor coding in a nostril-dependent manner and to correlate PC odor representations with subjects’ ability to identify odorants. We hypothesized that PC would temporally segregate odor information arising from the ipsilateral and contralateral nostrils and thus maintain distinct representations of odor identity for each nostril.

## RESULTS

### Experimental paradigm and behavioral performance

Ten subjects with intracranial depth electrodes participated in an odor identification task where odor stimuli were delivered either to the left, right, or both nostrils (bi-nostril condition), via a computer-controlled olfactometer (Figure 1A). For each odor type, the onset latency and amplitude of the odor pulses were matched between the left and right nostril conditions using a photoionization detector (PID) (Figure S1). Insertion of separate odor delivery tubes into each nostril (Figure 1A) ensured that odor delivery to one nostril did not contaminate the other, as validated with gas chromatography-mass spectrometry (Figure S2). Each trial began with a fixation cross, followed by a countdown and cue to sniff. Subjects received one of three unique odors (e.g, coffee, banana, eucalyptus) or odorless air through one of the three nostril configurations (Figure 1B, top panel). Subjects were subsequently asked to identify the stimulus they received in a four alternative forced choice task (odor identity judgment). In a subset of trials, subjects were asked a follow-up question on whether they received the stimulus from left, right, or both nostrils (laterality judgment). Air flow to both nostrils was constant and equal (4 L/min/nostril) throughout the experiment across all trial types. During stimulus delivery, valves for both left and right nostrils switched simultaneously from air to either a blank or odor line depending on the trial type (Figure 1B, middle and bottom panels). Local field potentials (LFPs) were recorded directly from the PC via intracranial depth electrodes throughout the task (Figure 1C).

**Figure 1.**
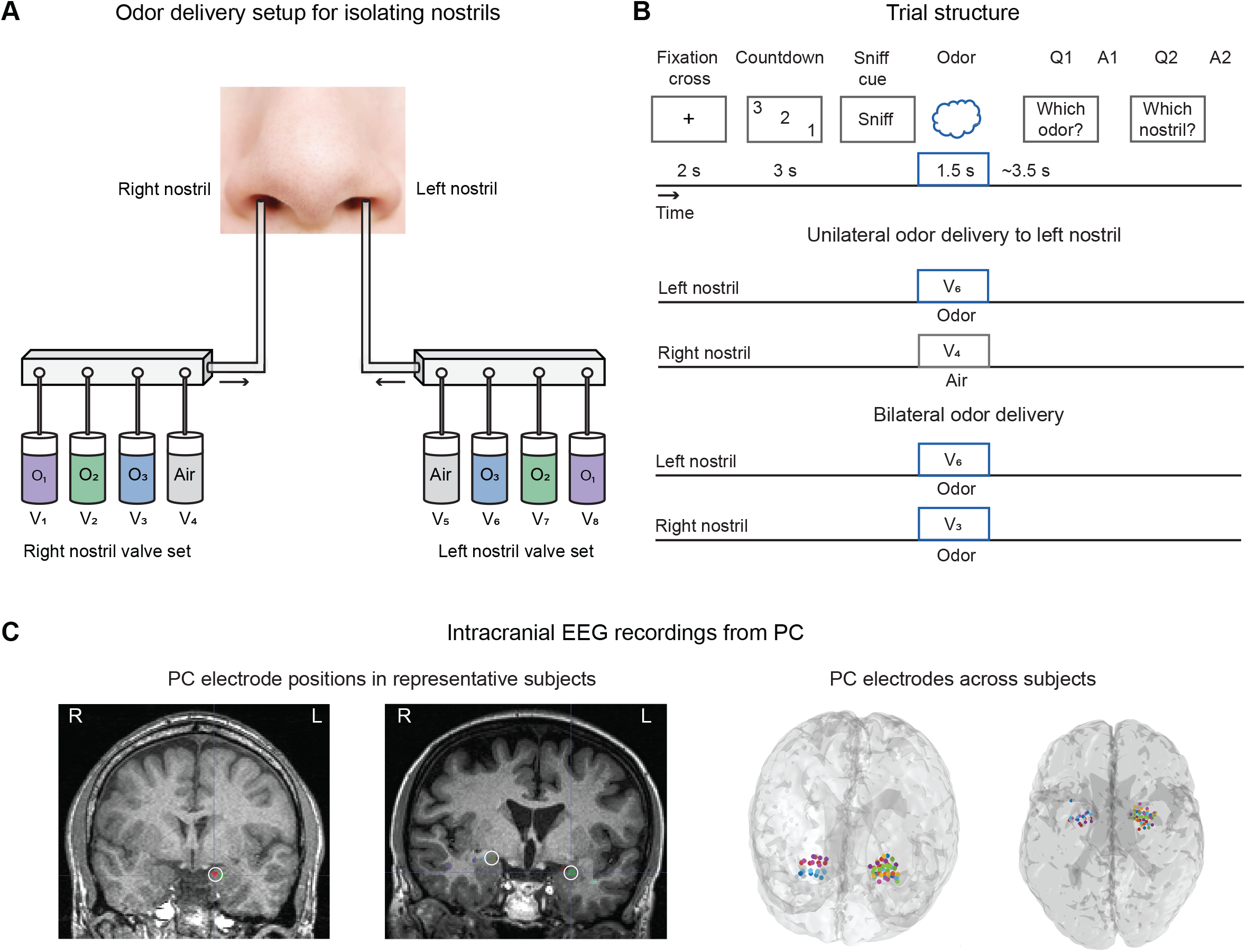
Experimental paradigm. **(A)** Isolating odor delivery to each nostril. Computer controlled olfactometer with dual mass flow controllers and valve sets enabled precise and isolated odor delivery to each nostril. Two small tubes were positioned ∼1 cm inside the nose and taped in place. The airflow to each nostril was kept constant at 4L/min throughout the entire experiment across all trial conditions. The odorants delivered to the left and right nostrils were matched in their onset latency, rise time and amplitude (see Figure S1) prior to each experimental session using a photoionization detector. This setup ensured odors delivered to one nostril did not contaminate the other, as validated with gas chromatography mass spectrometry (see Figure S2). **(B)** Trial and odor delivery structure. Top panel; Trials began with a fixation cross, followed by a countdown and cue to sniff. Subjects received one of three distinct odors or odorless air for 1.5 seconds in one of three nostril conditions (left, right or both nostrils). After each stimulus, subjects were asked to identify the odorant in a four alternative forced choice. In a subset of trials, subjects were also asked to identify which side the stimuli were delivered from in a three alternative forced choice question. Middle panel; schematic of uni-nostril odor delivery to the left nostril. Air flow is kept constant at 4L/min for both left and right nostrils. At stimulus onset, left nostril valve switches from odorless air to odor, while keeping the airflow constant, and the right nostril valve switches from odorless air to a second channel of odorless air, also maintaining the 4L/min airflow. This setup ensures that both left and right nostril valves switch on and off simultaneously and maintain the same airflow within each nostril. Bottom panel; bi-nostril odor delivery schematic illustrates that the valves for both nostrils switch to the same odor type simultaneously. The olfactometer setup ensured that onset latency and amplitude of odor delivery to left and right nostrils were matched (see Figure S1). **(C)** PC electrode locations. Left, position of PC electrodes shown on coronal MRI sections in two representative subjects with unilateral and bilateral PC electrodes, respectively. Each colored dot represents one contact point on a depth electrode and the contacts positioned in PC are circled in white. Right, all PC electrode contacts across subjects depicted on an average MNI-152 brain. Colors represent PC contacts from individual subjects.

A summary of behavioral results for each nostril condition is shown in Figure 2. In addition, we averaged across the left and right nostril trials to report direct comparisons between the uni-nostril versus bi-nostril conditions. Subjects correctly reported the presence of an odor (true positive rate) in 84.9% ± 2.5%, 89.3% ± 4.2%, and 97.8% ± 1.0% of left, right, and bi-nostril trials, respectively (Figure 2A, one-sample Wilcoxon signed rank test vs. 50% chance level, *p* = 0.002 for each nostril condition). Odor detection performance was higher for bi-nostril compared to uni-nostril trials, but there was no significant difference between the detection performance of left and right nostrils (two-sample Wilcoxon signed rank tests, p=0.002 and p=0.43, respectively).

**Figure 2.**
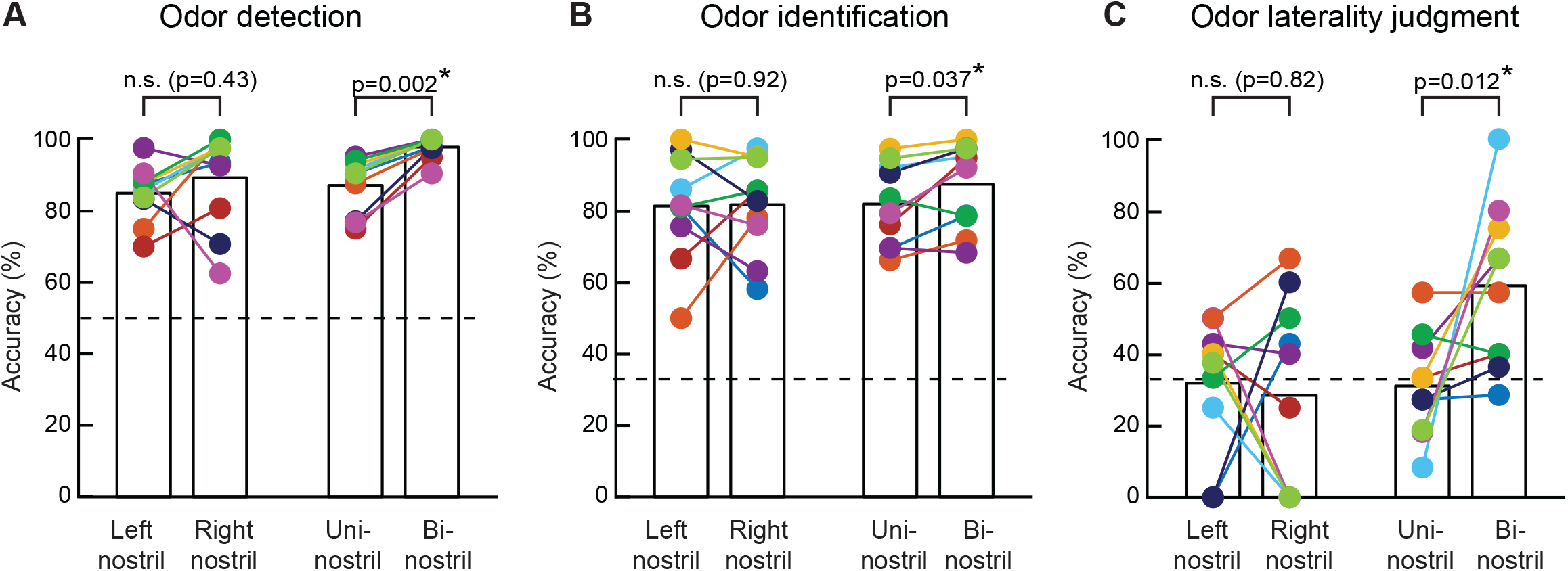
Behavioral performance. Task performance on odor detection **(A)**, odor identification **(B)**, and laterality judgment **(C)** for each nostril condition. Left and right nostril trials were averaged to illustrate behavioral performance in the uni-nostril conditions. In **(A-C)**, horizontal dashed lines indicate chance level accuracy of 50%, 33%, and 33%, respectively. Each colored circle indicates data from one subject. There was no difference between left and right nostril trials for any of the three behavioral metrics. There was a small but significant improvement in all three metrics for bi-nostril vs. uni-nostril trials (paired-sample Wilcoxon signed rank test, p-values shown on bar graphs). Not significant results shown as n.s. See also Figure S3.

For trials in which subjects correctly detected the presence of an odor, odor identification performance was 81.4% ± 4.7%, 81.7% ± 4.2%, and 87.4% ± 3.7% for left, right, and bi-nostril trials, respectively (Figure 2B, one-sample Wilcoxon signed rank test vs. 33.3% chance level, p=0.002 for each nostril condition). Whilst there was no significant difference in odor identification performance between the left versus right nostril conditions (two-sample Wilcoxon signed rank test, p=0.92), we observed a relatively small but significant improvement in identification of bi-nostril compared to uni-nostril odors (two-sample Wilcoxon signed rank test, p=0.037).

Finally, we asked if subjects could discern which nostril odor stimuli were delivered from (Figure 2C). Subjects could only designate bi-nostril trials above chance level (left nostril 31.9% ± 5.8%, right nostril 28.5% ± 8.5%, bi-nostril; 59.0% ± 7.2%, one-sample Wilcoxon signed rank test vs 33.3% chance level, p>0.05, p>0.05, and p=0.006, respectively). The laterality judgment was significantly better for bi-nostril versus uni-nostril trials but there were no significant differences between the left and right nostril trials (two-sample Wilcoxon signed rank tests, p=0.012 and p=0.82 respectively).

At the group level, there were no significant behavioral differences between left versus right nostril odor trials. We also confirmed these results at the level of individual subjects. We found that the majority of subjects did not display differences between the two nostrils. The number of subjects with a statistically significant difference was limited to 3/10 for odor detection (right nostril higher in two subjects), 2/10 for odor identification (right nostril higher in one), and 0/10 for odor laterality assessment (Fisher’s exact test, p<0.05).

Next, we asked if behavioral performance varied across odor types. For each subject, we compared odor detection, identification, and laterality assessment performance across the three odor types (odor A versus odor B, odor A versus odor C, odor B versus odor C), resulting in 30 total pairwise comparisons per behavioral metric. Behavioral performance was different only for a small number of odor pairs: 5 odor pairs for odor detection, 4 pairs for odor identification, and none for laterality assessment (Fisher’s exact test, p<0.05), suggesting that odor type did not play a strong role in driving behavioral effects.

Previous work suggests that nasal cycling (i.e., differences in air flow across the two nostrils) may lead subjects to sniff longer with one nostril over the other^26^. To assess if subjects sampled odors differently across nostril conditions we compared inhalation duration, latency to peak of inhalation, and exhalation duration. We did not observe differences in any one of the three sniff metrics between the left and right nostrils (Figure S3, paired-sample Wilcoxon signed rank test, p>0.05). Consistent with prior work^26^, subjects inhaled longer during uni-nostril versus bi-nostril odor trials (paired-sample Wilcoxon signed rank test, p=0.02).

Overall, our behavioral results show that subjects can detect and identify odors well above chance level in all nostril configurations, with the bi-nostril condition showing a relatively small but statistically-significant advantage in odor detection, odor identification and odor laterality assessment compared to the uni-nostril condition. Importantly, there were no differences in task performance and sniffing behavior between the left and right nostrils, suggesting that neither the left nor the right nostril conferred a relative advantage in odor sampling.

### Robustness of PC odor representations correlates with successful odor identification

We first sought to assess if odors elicit oscillations in PC and if the spectrotemporal characteristics of these responses vary across odor types. Single neuron studies in rodents and neuroimaging work in humans have shown that distinct odorants induce neural activity with distinct spatiotemporal profiles in PC^27–30^. In line with these observations, recent iEEG studies by our group have shown that PC oscillatory patterns also vary across odorants, and that differences in the frequency, power, and timing of odor-induced oscillations can be used to decode odor identity^31,32^.

To assess the presence of odor-induced oscillations, we aligned trials to sniff onset (time = 0 s) and conducted time-frequency analysis for each subject, averaging across odor types and nostril conditions. The grand-average time-frequency plot in Figure 3A shows the average piriform response across all subjects, odor types, and nostril conditions. Consistent with previously-published human PC recordings by our group^31,32^ and others^33,34^, there were robust odor-induced responses in the theta band (3-8 Hz) in the first second of the post-stimulus interval ([0-1]s, 4.04 ± 1.30 z relative to pre-stimulus baseline; one-sample Wilcoxon signed rank test vs. 0, *p* = 0.004), followed by responses in the beta/gamma frequency range (20-120 Hz) during the latter two seconds of the post-stimulus epoch ([1-3]s, 1.69 ± 0.59 z; *p* = 0.01).

**Figure 3.**
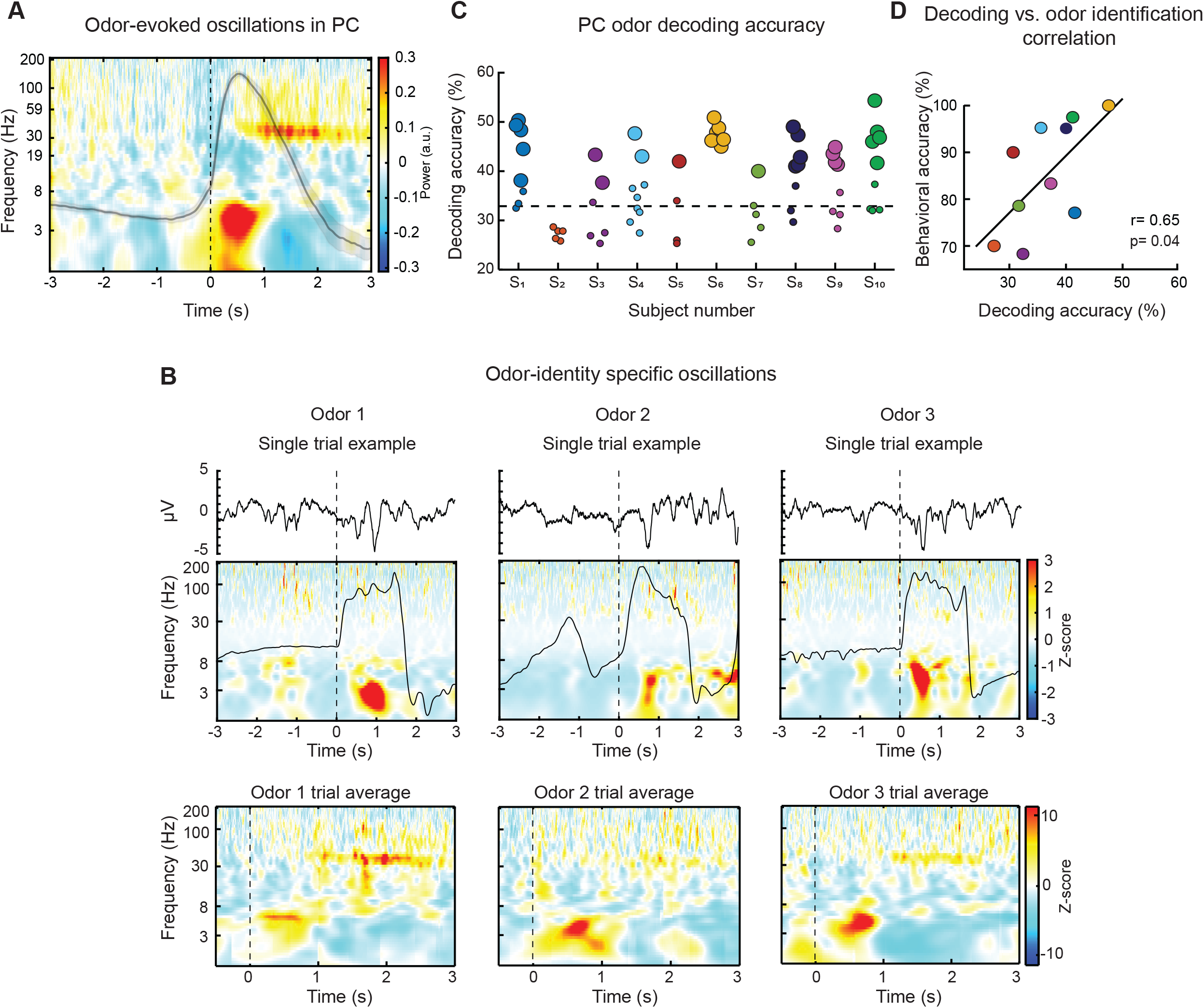
Behaviorally-relevant odor-induced oscillatory responses. **(A)** Odors induce robust oscillations in PC. Subject-averaged time-frequency plot depicting the grand-average odor response across all odor types and nostril conditions. Prior to averaging across subjects, subject-level power was first normalized with respect to the pre-stimulus baseline, and then normalized with respect to maximum power across time and frequency (see Methods). Plot is aligned to onset of inhalation (time = 0 s) during odor delivery. Average sniff trace is overlaid in gray (mean ± s.e.m. across subjects). **(B)** Distinct odor types induce distinct oscillatory dynamics. Top panel shows representative single-trial examples of local field potential traces and time-frequency plots from one subject (left column, citral, middle, coffee, right, eucalyptus). Power normalized with respect to the entire experimental trace. The bottom panel shows corresponding trial-averaged time-frequency plots for each of the three odor types. Power normalized with respect to pre-stimulus baseline. Note that the pattern of odor-induced oscillations across time and frequency varies for different odor types. **(C)** Odor identity could be classified with a model trained on time-frequency patterns of odor-induced responses. Decoding accuracy for bi-nostril odor trials is shown for each PC electrode across all subjects. Dashed line indicates theoretical chance level of 33%. Larger circles indicate PC electrodes with statistically significant decoding accuracy with respect to surrogate data obtained by shuffling the odor labels (one-tailed permutation test, p < 0.025; see Methods). **(D)** Greater disambiguation in PC odor representations correlates with better odor identification performance across subjects. Pearson’s correlation between decoding accuracy (averaged across all PC electrodes for each subject) and behavioral accuracy was computed across subjects using bi-nostril odor trials. Each colored dot indicates one subject, and solid black line shows linear fit.

Importantly, spectrotemporal features of odor-induced oscillations varied across odor types at the level of single trials. Representative single trial and trial-averaged spectrograms of one subject in response to three different odor types are shown in Figure 3B. To determine if the specific patterns of odor-induced oscillations at the single trial level contained information on odor identity, we implemented a three-way support vector machine (SVM) classifier for each PC electrode, following methods from our previous work^31,32^. Briefly, for each odor trial and PC electrode, we first extracted the spectrotemporal features from the 3-second post-stimulus interval. Then, for each PC electrode, we implemented a leave-out-one cross validation technique to train on all but one randomly-selected trial for each odor type, and tested classifier performance on the left-out set of trials. We sampled the minimum number of trials across odor types for the training set to prevent biasing the classifier by differences in sample size. Classifier performance for each PC electrode was quantified by repeating this procedure 200 times, and taking the average decoding accuracy across iterations. Statistical significance for each electrode was determined independently with respect to its own null distribution generated by randomly shuffling the odor labels (see Methods).

Following this approach, we first assessed which frequency band contained the most information about odor identity by training the classifier with either theta (3-8 Hz), beta/gamma (20-120 Hz), or broadband (3-120 Hz) oscillatory activity. Decoding accuracy was z-score normalized with respect to the corresponding null distributions in order to control for different number of features across the three frequency conditions. We found that the classifier trained on broadband oscillations performed significantly better than the classifier trained on theta oscillations alone (paired-sample Wilcoxon signed rank test, p=0.006) or beta/gamma oscillations alone (paired-sample Wilcoxon signed rank test, p=0.049) (Figure S4 C). We therefore performed all subsequent analyses with broadband oscillatory features. Using the subset of bi-nostril odor trials, we found that odor identity could be successfully decoded from broadband oscillations in at least one PC electrode in 9/10 subjects (decoding accuracy significant if >95^th^ percentile of null distribution, i.e., one-tailed permutation test, p < 0.025). The percentage of electrodes with significant decoding was 48.2% ± 9.0% across all subjects (Figure 3C, statistically significant electrodes shown as large circles). Next, we investigated if some odor types were more reliably decoded than others. Using the confusion matrices obtained from data presented in Figure 3C, we compared the sensitivity index (true positive/ true positive + false negative) across pairs of odor types in each subject (odor A vs. odor B, odor A vs. odor C, odor B vs. odor C). We found that none of the 30 odor pairs (across ten subjects) were significantly different from one another (Fisher’s exact test, p > 0.1).

As odor decoding accuracy was variable across subjects (Figure 3C), we examined whether this variation was systematically related to odor identification performance. We computed a Pearson’s correlation between odor identity decoding accuracy (for each subject, we took the mean decoding accuracy across PC electrodes) and odor identification performance across subjects, and found that the two variables were strongly correlated (Figure 3D, Pearson’s correlation, r=0.65, p=0.04).

Consistent with previous studies from our group^31,32^, we show that odor identity can be decoded from the spectrotemporal features of PC neural oscillations. Importantly, we provide novel evidence that enhanced disambiguation of PC odor representations correlates with improved odor identification performance.

### Ipsilateral odors are encoded faster than contralateral odors in PC

We next investigated if the speed and accuracy of odor coding in PC differs depending on which nostril odor information arises from. To assess the temporal evolution of odor coding in PC, we computed the decoding accuracy for each PC electrode in a time-resolved fashion, using 200-ms sliding windows with 50% overlap (see Methods). Trials were grouped into ipsilateral, contralateral, and bilateral conditions based on the site of odor delivery with respect to each PC electrode: for electrodes in the left hemisphere, left nostril odor trials were categorized as ipsilateral and right nostril odor trials were categorized as contralateral conditions and vice versa for the electrodes in the right hemisphere.

Importantly, to control for variability in inhalation length across subjects and uni-versus bi-nostril conditions (Figure S3 A, left panel; mean inhalation duration ranged between 1170 and 2662 ms across subjects), we normalized the time axis with respect to each subject’s mean inhalation duration in a given nostril condition such that 0% and 100% correspond to the onset and offset of inhalation, respectively. Limiting the analysis window to the inspiratory phase for each subject also ensured that our analysis was not confounded by potential re-entry of odorants into the nose upon exhalation, through the retronasal pathway^35^. Lastly, to control for differences in number of PC electrodes across subjects and to ensure that each subject contributed equally to the group results, we used a resampling procedure, in which we randomly sampled the minimum number of PC electrodes across subjects 200 times, and then took the average across iterations.

The subject-averaged results revealed striking differences in the relative time course of odor coding across the nostril conditions (Figure 4A). Uni-nostril odors arising from either the ipsilateral or the contralateral nostril were encoded within a single temporal epoch, though ipsilateral odors were encoded earlier than contralateral odors. Decoding of ipsilateral odors peaked at 40% (range: 32-47%; one-tailed permutation test, cluster-level p<0.025), whereas decoding of contralateral odors peaked later at 66% of inhalation (range: 58-73%, %; one-tailed permutation test, cluster-level p<0.025). Importantly, in contrast to uni-nostril odors, odor identity was encoded in two distinct, temporally-segregated epochs (i.e., dual coding) in the bi-nostril condition, with the first epoch peaking at 13% (range: 8-18%) and the second epoch peaking at 43% of inhalation (range: 37-57%). The statistical significance of temporal clusters were determined by first constructing null distributions for each nostril condition by randomly shifting the electrode-level time series prior to computing the group average. This was repeated 500 times, and the statistical significance was determined at the cluster level to account for multiple comparisons (see Methods). Notably, the temporal patterns observed at the group-average (Figure 4A) were also detectable at the single-electrode level (examples from three different subjects shown in Figure 4B). While the time course of odor coding varied depending on site of odor delivery, the peak decoding accuracy did not differ across the three nostril conditions (Figure 4C, paired-sample Wilcoxon signed rank test, p>0.05). In sum, odor identity was encoded just as robustly irrespective of the site of odor input, but odor identity information emerged faster for odors arising from the ipsilateral nostril compared to the contralateral nostril.

**Figure 4.**
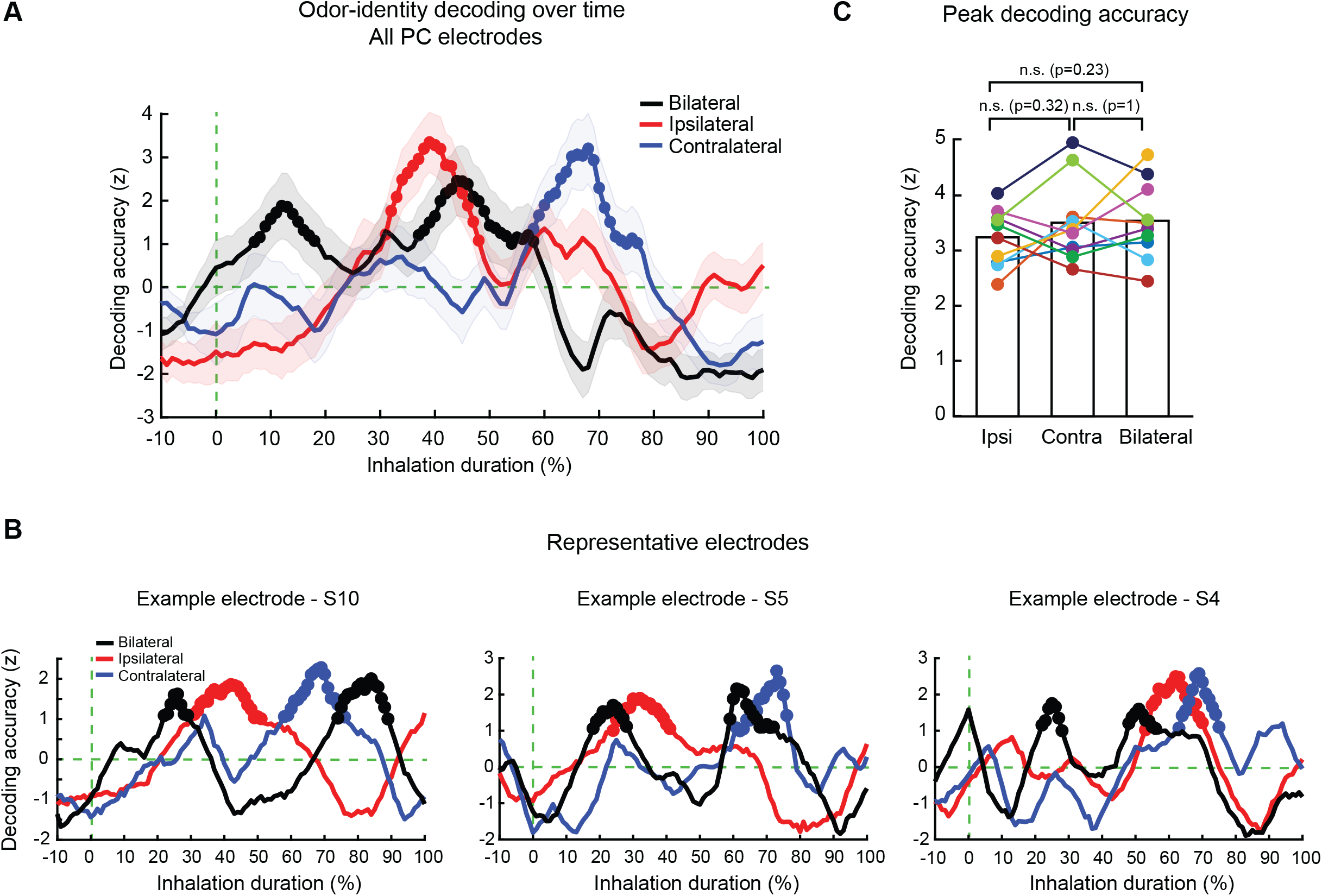
Ipsilateral and contralateral odors are encoded in temporally staggered single temporal epochs while bi-nostril odors are encoded in two distinct temporal epochs. (A) Subject-averaged time-resolved odor decoding analysis illustrates the differential time course of PC odor representations depending on stimulus laterality. Solid lines correspond to the mean and shaded lines represent the standard deviation of resampled distribution (see Methods). Time axis is shown as percentage of inhalation duration, where 0% and 100% represent the onset and end of inhalation, respectively. Decoding accuracy values were first normalized with respect to odor label shuffled surrogate data, and then normalized with respect to time-shifted surrogate data (see Methods). The latter null distribution was used to determine significant temporal clusters (one-tailed permutation test, cluster-level p<0.025), indicated with solid circles (see Methods). See also Figure S4 A-D. (B) Nostril-dependent temporal patterns of odor representations are also observed at the level of single electrodes. Each panel shows data from one PC electrode, obtained from three different subjects (S10, S5, S4). (C) The peak decoding accuracy throughout the inhalation period is not significantly different across nostril conditions at the group-level (paired-sample Wilcoxon signed rank test, p>0.05). Each colored dot represents one subject.

Next, we performed three additional analyses to confirm the above findings. First, to validate that our results were not an artifact of converting the time axis from seconds to percentage of inhalation, we repeated the same analysis in Figure 4A without normalizing the post-stimulus interval and found that the temporal pattern of odor coding remained qualitatively similar across both approaches (Figure S4 A).

Second, we conducted the same time-resolved decoding analysis in Figure 4A with just the theta band (3-8 Hz) or the beta/gamma band (20-120 Hz). Although these frequency bands contained less odor-identity information on their own (as shown in Figure S4 C), their temporal patterns were nonetheless qualitatively similar to broadband activity (Figure S4 D). Lastly, we asked whether the dual coding epochs observed in the bi-nostril condition may arise from higher variability in the timing of bi-nostril versus uni-nostril odor representations. To address this possibility we computed the variability in the timing of odor coding epochs either across electrodes (within each subject) or across subjects. First, we computed the variability in the peak time of the first significant decoding cluster across PC electrodes for each subject. We found that this inter-electrode temporal variability was lower in the bi-nostril compared to the ipsilateral condition (Figure S4 C, paired-sample Wilcoxon signed rank test, p=0.03) and not significantly different from the contralateral condition (paired-sample Wilcoxon signed rank test, p=0.5). Next we computed the median odor onset latency for each subject (median of the first significant decoding cluster across the subject’s PC electrodes) and computed the standard deviation of this onset latency metric across subjects. We found that the standard deviation of odor onset latency across subjects was comparable across the bilateral, ipsilateral and contralateral conditions (14.69, 15.89, and 21.72 respectively). These results suggest that the dual odor coding epochs in the bi-nostrils condition do not arise from increased variability in the timing of odor representations across electrodes or subjects.

Our findings suggest that when odors are presented to just one nostril, ipsilateral odors are encoded earlier than contralateral odors in PC. Interestingly, when the same odorant is presented to both nostrils simultaneously, PC odor representations emerge as two, temporally segregated clusters within the inhalation window.

### Ipsilateral and contralateral odors elicit similar yet distinguishable responses in PC

Our results show that ipsilateral odor representations in PC emerge earlier than their contralateral counterparts (Figure 4A and 4B). These findings raise the intriguing possibility that the same odor delivered to the ipsilateral versus contralateral nostril may ultimately elicit similar representations that are merely shifted in time. Indeed, work in rodents has shown that PC responses to ipsilateral versus contralateral odors are correlated at the neural population level^22^.

To test this hypothesis, we conducted time-frequency similarity analysis (see Methods) on ipsilateral and contralateral odor responses, enabling us to assess the overall similarity between odor representations across the two conditions. By training a classifier with single trials from one condition (e.g., ipsilateral) and testing the accuracy of the model using trials from the other condition (e.g., contralateral), we were able to quantify the similarity between the two conditions for each PC electrode in a time-resolved fashion. This analysis was conducted independently for all possible pairs of training and testing time bins tiling the inhalation period. The prediction here is that if the responses are similar to one another at a given pair of training and testing time points, then the classifier trained on one condition (e.g., ipsilateral) should successfully decode odor identity when tested with odor responses from the other condition (e.g., contralateral).

The resulting group-averaged time-frequency similarity matrices (TFSM) are shown in Figure 5. Values along the diagonal correspond to average decoding accuracy from training and testing at the same time points (e.g., train and test at 40% of inhalation), whereas the off-diagonal values show decoding accuracy from training and testing at different time points (e.g., train at 40%, and test at 80%). We also computed the group-average TFSM by reversing the training and testing conditions (i.e., train on contralateral and test on ipsilateral) to identify robust, bidirectional representational similarity (Figure 5B). Because we were interested in the representational similarity between two epochs that were temporally staggered (i.e., off-diagonal values in the TFSM), we computed the contrast between the two TFSMs in Figure 5A and 5B to eliminate the on-diagonal values (Figure 5C). The statistical significance of the resulting matrix was assessed via a surrogate data approach, where decoding accuracy values were circularly shifted across either training or testing time (see Methods). We found that ipsilateral responses occurring at 41-59% of inhalation could decode contralateral responses occurring later at 67-94% of inhalation, as shown by the positive cluster above the diagonal (two-tailed permutation test, cluster-level p < 0.05). Likewise, we observed congruent findings for the reverse direction: contralateral responses occurring at 67-88% of inhalation could decode ipsilateral representations at 41-61% of inhalation (negative cluster below the diagonal). These findings support our hypothesis that the same odor arising from the contralateral versus ipsilateral nostril generate similar but temporally-delayed representations in PC.

**Figure 5.**
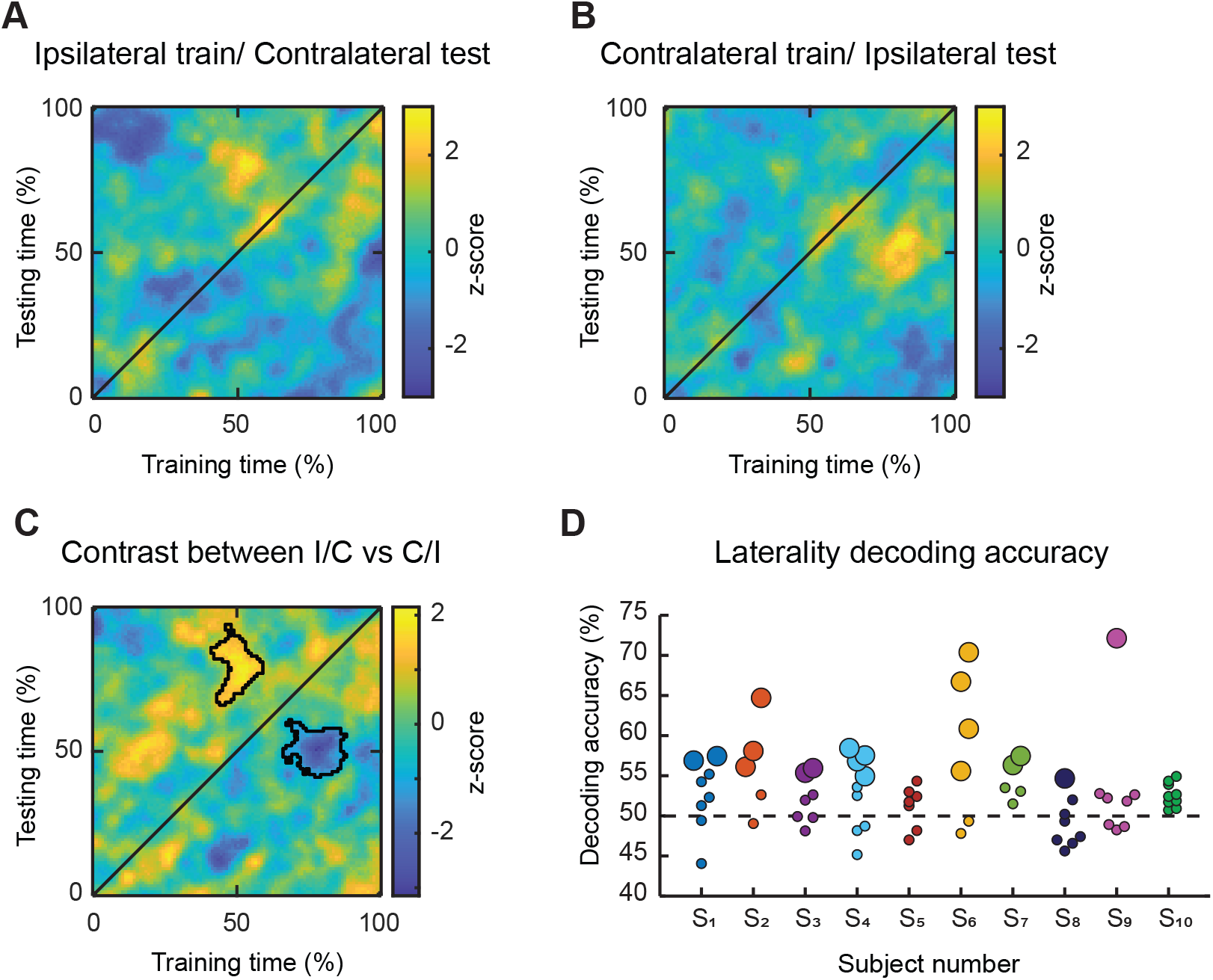
The same odorant delivered to ipsilateral versus contralateral nostrils induce similar yet distinguishable representations in PC. **(A-C)** Ipsilateral and contralateral odor representations were similar albeit shifted in time. Subject-averaged time-frequency similarity matrices (TFSM) constructed by training on ipsilateral and testing on contralateral odor responses **(A)**, or training on contralateral and testing on ipsilateral odor responses **(B),** across all pairs of time points tiling the inhalation duration. Values along the diagonal lines of TFSM show the subject-averaged decoding accuracy when the classifier was trained and tested with data from the same time points. Off diagonal values indicate when the classifier was trained and tested with data from different time points. Decoding accuracy values were normalized with respect to odor label shuffled surrogate data. The contrast between the subject-averaged TFSMs in **(A, B)** is shown in **(C)**, illustrating that early odor representations in the ipsilateral condition can be used to decode the later odor representations in the contralateral condition. Decoding accuracies in the contrast TFSM were further normalized with respect to surrogate data obtained by shuffling training and testing times which were used to identify significant clusters, as marked with black contours (two-tailed permutation test, cluster-level p<0.05; see Methods). Time axis is shown as percentage of inhalation duration (as in Figure 4). See Figure S3 for additional details. (D) Representations of ipsilateral and contralateral odors remained distinguishable. Time-frequency patterns of odor representations during the significant epochs (as outlined in the black contours in (C)) were used to decode odor laterality (ipsilateral or contralateral condition). Each circle shows the decoding accuracy of one PC electrode. The larger circles indicate statistical significance with respect to surrogate data obtained by shuffling the nostril labels (one-tailed permutation test, p < 0.025; see Methods). Dashed line indicates theoretical chance level of 50%.

Finally, we considered that the strong similarity between the ipsilateral and contralateral odor coding epochs does not preclude the possibility that these odor representations could nonetheless be differentiable on a fine-grained level. A two-way classifier was trained to decode stimulus laterality (ipsilateral versus contralateral) based on the spectrotemporal features of odor responses. To ensure that differences in the latency of odor representations would not influence classification results, we selected the time intervals with significant odor decoding for each nostril condition (see Methods). We found that responses to ipsilateral versus contralateral odors could be differentiated in at least one electrode in 7/10 patients (one-tailed permutation test, p < 0.025; Figure 5D). Across subjects, decoding was significant in 28.97% ± 7.44% of electrodes. Taken together, these results demonstrate that the same odor arising from the ipsilateral versus the contralateral nostril induces similar yet distinguishable representations that emerge at different intervals within the window of inhalation.

### Dual coding of bi-nostril odors arises from temporal segregation of ipsilateral and contralateral odor inputs

In contrast to uni-nostril odors, bi-nostril odors were encoded in two temporally-segregated epochs (i.e., dual coding; Figure 4A). Interestingly, the temporal distance between these two epochs closely matched the time lag between the ipsilateral and contralateral coding epochs in the uni-nostril conditions (29% versus 26% of the inhalation duration, respectively). Based on these observations, we asked if the two non-overlapping epochs observed in the bi-nostril condition correspond to representations of ipsilateral and contralateral odor inputs. We reasoned that if our hypothesis is true, then responses within the first coding epoch in the bi-nostril condition should resemble the responses within the ipsilateral coding epoch, whereas responses occurring in the second coding epoch should instead resemble the contralateral coding epoch. To assess the similarity of coding epochs across conditions we applied the time-frequency similarity analysis described above and compared odor coding in the bi-nostril condition with odor coding in the ipsilateral or contralateral conditions. Specifically, for each PC electrode, the classifier was trained on single trials using the bilateral odor responses and tested independently on ipsilateral or contralateral odor responses across all possible training and testing time points throughout inhalation. Similar to our approach in Figure 5, the training and testing sets were reversed for each comparison (Figure S5), and statistical significance was assessed on the resulting contrast matrices (two-tailed permutation test, cluster-level p<0.05). We found that bi-nostril responses at 12-34% of inhalation could decode ipsilateral responses at 43-65% of inhalation (Figure 6A). Moreover, bi-nostril responses occurring later at 39-69% of inhalation could decode contralateral responses at 57-93% of inhalation (Figure 6B). Importantly, the mirror-symmetric clusters above and below the diagonal lines indicate that reversing the training and testing conditions replicated the same results for both temporal epochs; ipsilateral responses at 41-65% could decode bilateral representations at 9-41%, and contralateral responses at 57-92% could decode bilateral representations at 21-65%.

**Figure 6.**
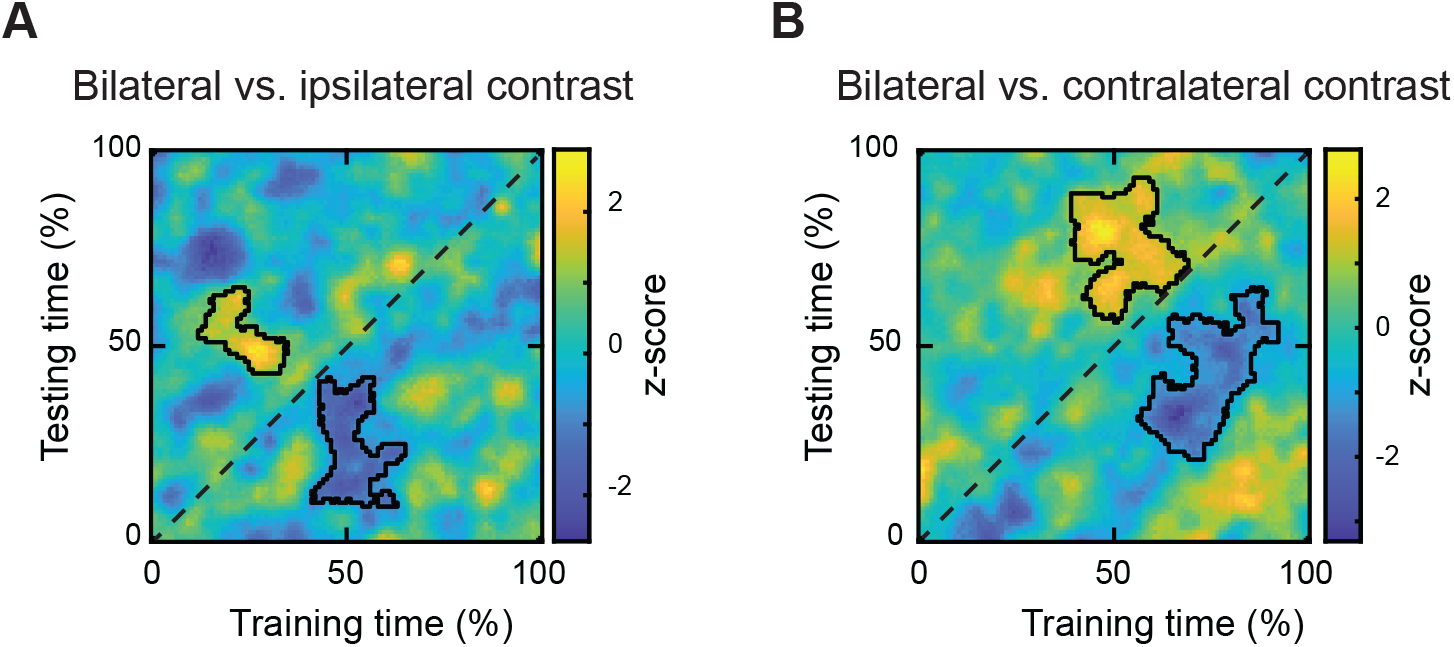
Dual coding of bi-nostril odors emerges from temporal segregation of ipsilateral and contralateral odor representations. Subject-averaged contrasts of time-frequency similarity matrices (TFSM) computed for bilateral vs. ipsilateral odor responses **(A)**, and for bilateral vs. contralateral odor responses **(B)**. Bilateral and ipsilateral odor representations were similar at an earlier time interval compared to bilateral and contralateral odor representations. Methods same as in Figure 5. See Figure S5 for additional details.

### Stimulus laterality dictates the time course of behaviorally-relevant odor coding

We next asked if the differences in the timing of odor representations (i.e., bilateral, ipsilateral, and contralateral) affected when PC odor representations inform odor identification performance. To address this question, we computed Pearson’s correlations between decoding accuracy and task performance across subjects in a time-resolved manner. This analysis is similar to the correlation shown in Figure 3D, except the correlations are computed independently at each time point for each laterality condition (see Figure S6 for methods schematic).

We found that decoding accuracy significantly correlated with odor identification performance for all three nostril conditions, but at distinct time points (Figure 7). To determine statistical significance, we implemented a surrogate data approach by shuffling decoding accuracy values across subjects, and computing correlations between the shuffled decoding accuracy values and odor identification accuracy values (see Methods). Significant correlations were observed at 33-47%, 49-66%, and 65-75% of the inhalation period for bilateral, ipsilateral, and contralateral conditions, respectively, with peak correlations occurring at 40%, 54%, and 70% of inhalation (one-tailed permutation test, cluster-level p<0.025). The temporal order of behaviorally-relevant odor coding recapitulated the order in which odor representations emerged across nostril conditions. Interestingly, in all three conditions, correlations with behavior arose later during the inhalation period compared to the onset of the corresponding odor coding epochs. For instance, bi-nostril odor representations emerged as early as 8% of inhalation but significant correlations with odor identification performance arose at 33% of inhalation. Similarly, the onset of significant correlations were delayed by 17% and 7% of the inhalation duration for ipsilateral and contralateral conditions, respectively. Lastly, while bilateral odors were encoded in two non-overlapping temporal clusters (occurring at 8-18% and 37-57% of inhalation, as shown in Figure 4A), correlations with behavior occurred within a single temporal cluster at 33-47% of inhalation. Together these data underscore the idea that just as stimulus laterality dictates the speed of odor coding in PC, stimulus laterality also affects how rapidly PC odor representations are able to inform odor identification.

**Figure 7.**
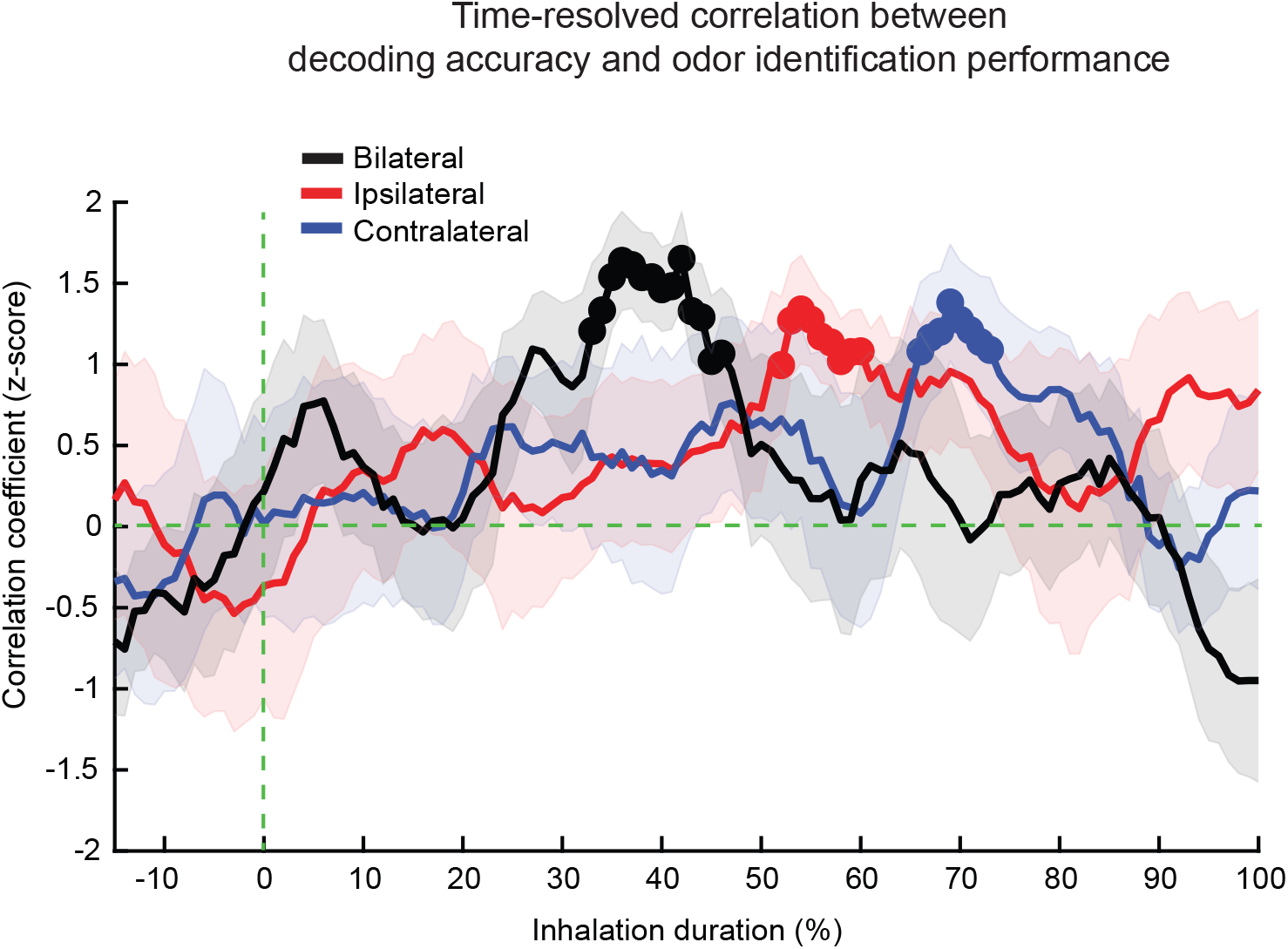
Time course of behaviorally-relevant odor coding depends on stimulus laterality. PC odor representations predicted behavioral performance at different time points depending on stimulus laterality. Subject-averaged time-resolved correlations between odor decoding accuracy and odor identification performance. At each time point, a Pearson’s correlation was computed between PC decoding accuracy and behavioral accuracy across subjects (as in Figure 3D). Correlation coefficients were normalized with respect to null distributions obtained by shuffling behavioral accuracy values across subjects (see Methods). These null distributions were also used to identify significant temporal clusters (shown as solid circles, one-tailed permutation test, cluster-level p<0.025). Solid lines and shaded areas represent the mean and standard deviation of the resampled distribution (see Methods). See also Figure S6.

## DISCUSSION

One key feature that binds the olfactory systems of all mammals, including humans, is the presence of two nostrils, each conveying a potentially unique snapshot of the external olfactory world. Here, using iEEG recordings and machine learning methods, we show that odor information from the two nostrils is temporally segregated in the primary olfactory cortex of humans. Furthermore, we demonstrate that greater differentiation among PC odor representations scales with improved odor identification across subjects. Lastly, we show that the nostril-dependent order of odor representations influences when PC odor informs odor identification.

### Temporal segregation of ipsilateral and contralateral odor information in PC

In the uni-nostril condition, we found that ipsilateral odors are encoded earlier than contralateral odors by 26% of the inhalation duration, which corresponds to 478 ms on average. Although ipsilateral and contralateral odors were encoded at different phases of inhalation, we found that stimulating either nostril with the same odor elicited similar yet distinguishable representations during their respective coding epochs. Early work in anesthetized rats has shown that odor responses in PC can be nostril-specific^21^, suggesting that a given odorant may activate different neurons in PC depending on stimulus laterality. Recent work in awake mice has found that neural population responses to ipsilateral versus contralateral odorants were indeed distinguishable but remained highly correlated^22^. Our results are consistent with previous work in rodents, and taken together suggest that ipsilateral and contralateral odors activate partially overlapping but distinct ensembles of neurons that elicit similar yet distinguishable response dynamics at the population level.

Interestingly, the temporal distance between the onset of ipsilateral and contralateral coding epochs in the uni-nostril condition (26% of the inhalation duration) was preserved when the same odorant was delivered to both nostrils simultaneously (29% of the inhalation duration, and 498 ms on average). One possible interpretation is that bi-nostril sniffing provides PC with temporally individuated snapshots of odor information from each nostril within one sniff. An alternative explanation is that at the single-trial level, the PC selectively encodes either the ipsilateral odor input (earlier during inhalation) or the contralateral odor input (later during inhalation), but not both. In this alternate scenario, odor input from each nostril is still temporally segregated into distinct segment of the inhalation cycle, albeit the dual coding epochs would only be observed in trial-averaged data. Future studies should investigate single-trial dynamics to distinguish between these two alternative scenarios. While the encoding of bi-nostril odors recapitulated the temporal pattern expected from taking the sum of the ipsilateral and contralateral patterns, there was one important difference: in the bi-nostril condition, epochs corresponding to ipsilateral and contralateral nostrils emerged earlier than their uni-nostril counterparts by 24% and 21% of the inhalation duration, respectively. Why simultaneous stimulation of both nostrils gives rise to faster odor coding than stimulation of just one nostril remains unclear. One marked difference of course is that the subjects received twice the amount of odorant in the bi-nostril versus uni-nostril trials. As such, it is possible that the faster emergence of odor representations may be explained by either the increase in total amount of odorant, and/or by computational advantages that bi-nostril smelling may confer to odor processing. Future experiments where subjects receive the same amount of total odor in the uni-nostril and bi-nostril conditions would provide a crucial control to address this question.

It is important to note that the temporal delay between ipsilateral and contralateral odor representations is not driven by differences in odor input across the two nostrils. For each odor type and experimental session, stimuli delivered to the left and right nostrils were calibrated using a photo-ionization detector to ensure that stimuli were matched in concentration, onset latency as shown in Figure S1. Furthermore, since there were no external cues indicating the site of odor delivery in a given trial, and the subjects received identical air flow in both nostrils throughout all trials, the observed results cannot be explained by potential sensory or anticipatory cues biasing attention to either nostril and altering odor coding in a nostril-specific fashion. Finally, an additional factor to consider is nasal cycling. Previous work has shown that one nostril typically has greater airflow than the other,^36,37^ and this may lead subjects to take longer sniffs when smelling through the nostril with the lower flow rate^26^. While we did not directly measure nasal cycling in our experiment, we did not observe such behavioral differences indicative of nasal cycling in our data. First, there were no significant differences in how well subjects were able to detect, identify, or lateralize odorants between the left versus right nostrils, and second, there were no significant differences in sniff metrics between the two nostrils. Therefore, our data suggest that putative nasal cycling did not bias subjects’ odor perception and sniff behavior to one nostril. Lastly, our analyses grouped trials into ipsilateral and contralateral conditions such that each condition contained data from both the left and the right nostril odor trials. As such, any putative advantages one nostril may have had would be reflected equally in both the ipsilateral and the contralateral conditions.

While our experiment cannot speak directly to the mechanisms underlying the temporal difference between ipsilateral and contralateral odor coding, recent studies in rodents suggest that information from ipsilateral versus contralateral nostrils may reach PC through distinct pathways (as summarized in Dalal et. al.^16^). Ipsilateral inputs are just two synapses removed from PC, as olfactory sensory neurons project to the ipsilateral olfactory bulb, which then project directly to the ipsilateral PC. Contralateral inputs, on the other hand, can follow one of two pathways to reach PC, via the so-called direct and indirect pathways, each requiring three or more synapses between the contralateral nostril and PC^16,19,20^. These differences may potentially explain why ipsilateral odor representations emerge faster in PC. In fact, early work using non-invasive scalp recordings in humans have shown that responses to ipsilateral odors arise faster compared to contralateral odors in the ipsilateral hemisphere^38–40^. However, these studies focused on the onset of odor-evoked responses (i.e., when neural activity changes from baseline), as opposed to onset of odor representations (i.e., the first time point in which odor identity can be decoded from neural activity), and furthermore did not investigate how these temporal differences influenced sensory convergence in the bi-nostril condition.

### Odor coding latency dictates time-course of behavioral correlations

PC is thought to be a critical region in encoding odor identity information and informing odor object perception^10^. Perhaps not surprisingly then, we observed that better disambiguation of odor representations in PC correlated with improved odor identification performance across subjects. Furthermore, the temporal order of time-resolved correlations between odor coding and behavioral performance mimicked the order of odor representations across nostril conditions: bi-nostril odor coding informed odor identification first, followed by ipsilateral, and lastly by contralateral odors. However, the time course of correlations between odor coding and odor identification was different than the time course of odor representations *per se*, in two important ways. First, for all three conditions, the behavioral correlations emerged later than the corresponding encoding epoch and second, in the bi-nostril condition, there was only a single epoch in which odor coding correlated with behavior. These observations suggest that PC odor representations influence downstream computations and odor identification with a time lag of 16% of the inhalation duration (corresponding on average to 275 ms) and furthermore imply that even though bi-nostril odors are encoded in two distinct epochs, these odor codes may nonetheless inform behavior within a singular epoch.

Altogether, our findings have important implications for odor coding in the olfactory system and provide evidence that human PC maintains distinct representations of odor information arising from each nostril through temporal segregation. If PC can temporally segregate identical odor inputs that arrive simultaneously to both nostrils, how then do naturally occurring differences in the timing, concentration or identity of odor inputs across the two nostrils affect odor coding in PC? Finally, these results raise the question of whether the human olfactory system, akin to the auditory system using interaural time differences to localize sounds^41^, can engage such a coding scheme to compare odor inputs across nostrils and aid in rapid odor localization within a single sniff.

## ACKNOWLEDGMENTS

The authors thank the patients who participated in this experiment and the staff at the Hospital of University of Pennsylvania Epilepsy Monitoring Unit for their assistance. We also thank Dr. Brianne Linne and Dr. Aline Robert-Hazotte for their assistance with the GC-MS. This work was supported by NIDCD grant R01DC018075 (J.A.G) and NINDS Kirschtein NRSA T32 postdoctoral grant (A.I.Y).

## AUTHOR CONTRIBUTIONS

Conceptualization, G.N.D, K.A.D, J.A.G; data collection, G.N.D, N.S; methodology, G.N.D, A.I.Y, N.S; data analysis, A.I.Y and G.N.D; resources, T.L, H.I.C, K.A.D, J.A.G; writing, G.N.D, A.I.Y, J.A.G; visualization, G.N.D, A.I.Y; supervision, J.A.G.

## STAR Methods

## RESOURCE AVALIABILITY

### Lead Contact

Further information and requests for resources should be directed to and will be fulfilled by the lead contact, G. Nazli Dikecligil

### Materials Availability

This study did not generate new unique reagents.

### Data and Code Availability

De-identified intracranial EEG data has been deposited at iEEG.org and are publicly available at the time of publication. This paper does not report original code. Additional information required to reanalyze the data reported in this paper is available from the lead contact upon request.

## EXPERIMENTAL MODEL AND SUBJECT DETAILS

Intracranial EEG recordings from human PC were obtained from ten subjects (S1-10, 2 male, mean age 30.9 years, range 23-47 years) with drug-resistant epilepsy who were implanted with depth electrodes for pre-surgical localization of seizure foci. Data were recorded from the Hospital of the University of Pennsylvania. Informed consent was obtained from all subjects in accordance with the local Institutional Review Boards.

## METHOD DETAILS

### Data acquisition

Electrode placement decisions were made exclusively on clinical grounds, and typically focused on areas in the temporal lobe. Overall, five subjects had bilateral piriform cortex (PC) electrodes, one subject had right-sided electrodes only, and four subjects had left-sided electrodes only. Implanted depth macro-electrodes (Adtech AG, Frankfurt, Germany; Model SD10R-SP05X-000) were of the following specifications: each lead had 8 or 12 cylindrical platinum electrode contacts, with 2.4 mm of exposed surface, and a 2.6 mm gap between contacts. Data were acquired using a 256-channel Natus recording system (Natus Medical Incorporated, Pleasanton, CA). For each subject, signals were sampled between 500 and 2000 Hz at the physician’s discretion. Respiratory airflow data were simultaneously collected using a piezoelectric pressure sensor attached to a nasal cannula (Salter Labs, Vista, CA).

### Electrode selection

After co-registering the pre-operative high-resolution MRI with the post-operative CT using a linear affine registration^42^, electrodes were localized in native space, and then transformed to Montreal Neurological Institute (MNI) space for visualization across all subjects on an average MNI-152 brain. Electrodes in PC were then determined based on visual comparison with manually-drawn regions of interest in MNI space. The number of PC electrodes across subjects were 7.40 ± 0.50 [5, 9] (mean ± s.e.m. [min, max]).

### Behavioral paradigm

We devised an olfactory task in which odor stimuli were presented through one of three stimulus routes (laterality): left nostril, right nostril, or both nostrils (bi-nostril). Odor trials were interspersed with trials in which odorless air was delivered to both nostrils. The subjects received continuous and steady air flow to each nostril throughout the entire experiment (4 liters per minute per nostril). During stimulus delivery (where one of 3 odorants or odorless air was presented), the valves for both nostrils switched simultaneously to either an odor line or to a control odorless airline depending on the trial condition (Figure 1B middle and bottom panels). The flow rate was computer controlled with mass flow controllers (MFCs, Alicat Scientific). This design ensured that (1) total airflow to the subject was constant throughout the entire experiment across all trials, (2) airflow to each nostril was identical across all trials irrespective of whether the nostril was receiving odor or odorless air, and (3) valves for both nostrils switched on and off simultaneously in each trial. In each odor trial, the delivered odor was selected pseudo-randomly from a set of three unique and discriminable odor types, which was delivered via a stimulus route that was also pseudo-randomly selected. There were no sensory cues indicative of the upcoming odor type and nostril condition. For each subject, the three odor types utilized for the entire experiment were selected from the following set: banana, coffee, orange, peanut butter, limonene, pine, and eucalyptus. Odor sets were different across subjects to ensure the results were generalizable to many odorants and were not driven by odorant-specific effects. Stimuli were administered via a custom-built, twelve-channel olfactometer. The olfactometer, equipped with two mass flow controllers (Alicat Scientific), and two sets of solenoid valves with 6 valves each (NResearch Inc.), was controlled using a computer equipped with MATLAB (MathWorks, Natick, MA), and the experiment was designed and administered using the PsychToolbox package^43^. For each odor type, the concentration and onset latency of the odorant delivered to the left and right nostril was matched. Figure S1 A illustrates the temporal profiles of the odor types used in the experiment: as shown in the insets and quantified in Figure S1 B, the odor onset latency, odor rise time, and area under the odor curves were matched between the left versus right nostril delivery for each odor type. Prior to each experiment, the odors were tested and calibrated with a photo-ionization detector (miniPID, Aurora Scientific) to ensure that the onset latency and peak intensity of odorants were matched for each odor type across the left and right nostril delivery.

Each trial began with a fixation cross, followed by a three second countdown. A visual cue instructing the subjects to take a sniff appeared after the countdown. This baseline period between the fixation cross and sniff cue lasted five seconds, and the stimulus duration was 1.5 seconds. During the experiment, each subject underwent a total of approximately 170 trials, and 15-25% of all trials were odorless air trials. Of the remaining trials, each unique odor type and stimulus route occurred in equal proportions.

After stimulus delivery, subjects performed up to two judgments, the first of which was presented 3.25-4 seconds after odor onset. First, subjects were provided with the labels of all three odor types as well as the option of “no odor,” and asked to identify the stimulus (across subjects, response time [RT] 3.92 ± 0.72 s). Second, for a pseudo-randomly selected subset of trials (approximately 30 trials across both odor and odorless air trials), subjects then reported whether the stimulus was delivered to the left, right, or both nostrils (RT 7.91 ± 2.54 s). Note that the subjects could only provide their answers after the question appeared on the monitor, as opposed to being able to provide a response as soon as they sampled and identified the odorant.

Behavioral performance was assessed for odor detection, odor identification, and stimulus laterality. As the neural data analysis was performed exclusively on odor trials, odor detection accuracy was calculated for the subset of trials in which subjects received an odor (i.e., true positive rate, chance-level detection accuracy is 50%. We note that for odorless air trials, the correct rejection rate was 79.09 ± 6.49 (mean ± s.e.m.) across all ten subjects (7/10 subjects had correct rejection rates above 75%).

Identification accuracy was computed for the subset of odor trials in which the presence of an odor was not missed (e.g., did not answer “no odor” when orange was presented). Hence, incorrect answers were those in which the delivered odor was detected, but misclassified (e.g., answer “coffee” when pine was presented), resulting in a chance-level accuracy of 33.3%. Similarly, to determine laterality accuracy, we calculated the percentage of correct responses in trials where the subject correctly detected the presence of an odor (chance-level accuracy 33%).

### Data pre-processing

All data were analyzed offline in MATLAB using custom scripts in conjunction with functions from the EEGLAB toolbox^44^ and other publicly available toolboxes^45,46^. Data were down-sampled to 500 Hz (after an anti-alias filter), and then high-pass filtered at 0.1 Hz using a two-way least-squares zero phase-lag finite impulse response (FIR) filter (eegfilt.m). Data were then notch-filtered at 60 Hz and its harmonics to remove line noise using a second-order infinite-impulse response (IIR) filter. We visually inspected all channels to eliminate channels with significant interictal epileptiform discharges. All extra-parenchymal electrodes were also eliminated based on visual inspection of electrodes in each subject’s native MRI space. Finally, we rejected noisy and flat channels in which the mean magnitude of the signal (mean of the absolute value of raw amplitudes) was 3 s.d. above or below the mean (across all included intracranial electrodes), respectively. Following our previous studies^31,32^, each remaining electrode’s LFP signal was then referenced to the common average (i.e., average signal across all included intracranial electrodes) to reduce common noise from, e.g., volume conduction and remote field effects^47^.

Further artifact rejection was conducted at the single trial level using a previously-reported event-level algorithm^48^, which was designed to automatically identify artifacts such as eye-blink artifacts, sharp transients, and interictal epileptiform discharges. Specifically, for each time point, we calculated the following metrics: 1) absolute amplitude; 2) gradient (first derivative); and 3) amplitude of the data following a 150 Hz high-pass filter. Each of these metrics were then independently *z*-score normalized with respect to the electrode-specific mean and s.d. Time points in which any of these measures exceeded 5 s.d., or all three measures simultaneously exceeded 3 s.d. were marked as artifacts. Data from 100 ms before and after each of these time points were then classified as artifacts and eliminated from all analyses.

Data were aligned to the onset of sniff in the presence of odor stimuli. As the identification question was presented as early as 3.25 seconds following stimulus onset across subjects, data analysis was limited to 3 seconds following stimulus presentation. An equivalent-length baseline period was extracted from each trial ([-3, 0] s).

Respiratory data were low-pass filtered at 10 Hz and manually inspected. Odor stimulus trials in which inspiration was sub-optimal were rejected if any of the following conditions were met: 1) overlap between odor and inhalation <100 ms; 2) peak velocity of inspiration <5^th^ percentile of the distribution of peak velocities across all odor trials (within each subject); 3) two successive rapid sniffs taken during the odor delivery period.

## QUANTIFICATION AND STATISTICAL ANALYSIS

All analyses were limited to odor trials, excluding odorless air trials. All group-level results were based on one data point from each subject, taking the mean across PC electrodes within subjects, unless otherwise noted. Finally, all statistical tests were two-tailed with α = 0.05 or one-tailed with a corrected α.

### Oscillatory power

For spectral analysis across time and frequency, raw traces were filtered between 1 and 200 Hz in 60 steps with center frequencies increasing logarithmically using a zero phase-lag FIR filter with 50% overlapping bandwidths. Subsequently, the Hilbert transform was applied to extract instantaneous amplitude/power at each time point ([-3, 3] s) and frequency bin. For depiction of single trial time-frequency (TF) results (Figure 3B, middle row), oscillatory power was *z*-score normalized with respect to the entire experimental recording within each frequency sub-band. For electrode-level or subject-level trial-averaged TF results (e.g., Fig 3B, bottom row) oscillatory power was *z*-score normalized with respect to the baseline period ([−3, −0.5]) within each frequency sub-band. To obtain subject-level TF results, we resampled the minimum number of trials across electrodes 200 times (to control for variable trial numbers after artifact rejection). Subject-level power in each frequency range of interest (theta [3-8 Hz] or beta/gamma [20-120 Hz]) was quantified by selecting the frequency sub-band (10 of 60 sub-bands spanned theta, 20 sub-bands spanned beta/gamma) with the most robust odor-induced activity within each frequency range, and then taking the mean across time within the corresponding post-stimulus epoch of interest. Finally, for group-level results (Figure 3A), subject-level oscillatory power in each sub-band was normalized a second time by dividing spectral power values in each TF pixel by the maximum power value within the post-stimulus prior to taking the average across subjects.

### Decoding odor identity from odor-induced patterns of oscillatory activity

We performed classification of odor identity from single-trial TF patterns of odor-induced oscillatory activity using a multi-class (three-way) non-linear support vector machine (SVM) with a radial basis function kernel^46^. As odors were randomly selected from a set of three unique odor types, chance-level accuracy is 33.3%. Following our previous studies^31,32^, we developed classifier models independently for each electrode.

To extract TF features, we used the same parameters as in the TF plots: baseline-normalized mean power in each frequency sub-band spanning the frequency range of interest (40 sub-bands spanning 3-120 Hz) was calculated across non-overlapping 10 ms windows tiling the post-stimulus epoch ([0, 3] s). To reduce computational cost, we averaged over contiguous frequency sub-bands (resulting in 20 sub-bands). For decoding with theta-band or beta/gamma-band oscillations only, TF features were calculated across 10 sub-bands spanning 3-8Hz, or 20 sub-bands spanning 20-120 Hz, respectively.

Model performance for each electrode was quantified using a leave-one-out cross-validation technique to train on all but one random set of trials from each of the three odor type classes, and tested on the left-out set of trials. We repeated this procedure 200 times and reported the mean of the resulting distribution of decoding accuracies. At each iteration, the minimum number of trials across odor types was sampled for the training set to ensure that the classifier would not be biased by differences in sample size. Statistical significance was determined using a non-parametric surrogate data approach. To do this, we constructed independent null distributions for each electrode (N = 500), whereby at each iteration, the classifier was built with shuffled labels, and then used to decode for the shuffled labels in the left-out set of trials. Importantly, in generating surrogate data, the feature space for each trial was not altered. Electrode-level significance was observed if the true decoding accuracy was in the top 2.5^th^ percentile of the null distribution.

We used the subset of bi-nostril odor trials to show model performance for each PC electrode across all subjects (Figure 3C). We also used the sub-set of bi-nostril odor trials for analysis of the correlation between odor identity decoding accuracy from the entire post-stimulus epoch and behavioral performance on the odor identity judgment (Figure 3D). Specifically, a Pearson’s correlation was performed using the raw decoding accuracy and the raw behavioral accuracy. We note that the results were qualitatively similar when the Pearson’s correlation was computed after *z*-score normalizing decoding accuracies with respect to null distributions constructed by label shuffling (as described above), and after independently *z*-score normalizing both decoding and behavioral accuracies across subjects to achieve a normal distribution.

### Time course of odor coding across stimulus laterality conditions

To investigate the time course of odor coding, we performed the same odor identity decoding analysis as outlined above, but in a time-resolved fashion. Features were extracted separately from 200 ms sliding windows (400 features: 20 time bins × 20 frequency bins) spanning −0.5 to 3 seconds (with respect to sniff onset) with 50% overlap, resulting in 36 decoding time bins. The time course of odor coding was compared across the following stimulus laterality conditions: 1) ipsilateral (odor delivered from ipsilateral nostril), 2) contralateral, and 3) bi-nostril. Note that in subjects with bilateral PC coverage, model performance was averaged across both left PC electrodes during left nostril stimulation and right PC electrodes during right nostril stimulation for the ipsilateral condition. Similarly, for the contralateral condition, model performance was averaged across left PC electrodes during right nostril stimulation and right PC electrodes during left nostril stimulation. As described in the section above (**Decoding odor identity from odor-induced patterns of oscillatory activity)**, surrogate data was generated for each electrode × decoding time bin by shuffling odor identity labels across trials (N = 500). This distribution was used to *z*-score normalize the observed decoding accuracy values. We then computed the group-level time series of decoding for each of the three stimulus laterality conditions. To do this, for each decoding time bin, we resampled the minimum number of PC electrodes across subjects 200 times to control for variable electrode numbers across subjects, which were then averaged. We then applied a one-tailed threshold of 1 s.d. to identify all temporally contiguous suprathreshold clusters.

To determine the statistical significance of the detected temporal clusters, we implemented a cluster-level approach that is widely used in the EEG literature^49^. Each suprathreshold cluster was represented by the sum of the z-statistics across its contiguous time bins, referred to as its cluster-level *z*-statistic. We then constructed a null distribution (N = 500) of cluster-level *z*-statistics using a surrogate data approach. Specifically, at each iteration, we repeated the same resampling procedure as above, except we circularly shifted each electrode-level decoding accuracy time series by a random amount, and then took the average across resampled electrodes across all subjects. At each iteration, we used the same *z*-score threshold of 1 z at each time bin on the average time series, and then took the cluster-level *z*-statistic of the largest suprathreshold cluster. This procedure was performed independently for each of the three laterality conditions. Suprathreshold clusters observed in each group-level time series were deemed statistically significant if they were in the top 2.5^th^ percentile of the corresponding null distribution. For depiction of group-level results (Figures 4A and S4 A), we *z*-score normalized values in each group-level time series with respect to the corresponding null distribution.

For single-electrode results (Figure 4B), we followed the same procedure as detailed above with the following difference: as there was only one electrode-level time series, the null distribution was generated by shuffling decoding accuracy values across time points (as opposed to circularly shifting the time series). Significant clusters identified at the single-electrode level were used to compare the variability of odor representation onsets across stimulus laterality conditions (Figure S4 B).

Finally, to control for inter-subject variability in sniff duration, significant clusters were also identified after normalizing time with respect to subject-specific inhalation durations, i.e., the duration from sniff onset until the end of the inhalation period (across subjects, 1816 ± 136 ms [1170, 2662]). To do this, subject-level decoding time courses were interpolated across 100 time points that spanned each subject’s mean inhalation duration (across trials) from 0 to 100%. We normalized time with respect to the mean inhalation duration (across trials) separately for the bi-nostril condition and for the two uni-nostril conditions.

### Similarity of odor representations across stimulus laterality conditions

Classifier models trained with odor trials in which odors were delivered through the ipsilateral, contralateral, or bi-nostril routes were then tested on odor trials from an alternate stimulus route.

Specifically, the classifier was trained with data from odor trials in which the stimulus was delivered from the ipsilateral or contralateral nostril, and then tested on patterns obtained from odors delivered from the contralateral or ipsilateral nostril, respectively (Il→Cl, Cl→Il; Figure 5A, B). In addition, we trained with data from bilateral odor stimulation, and tested on patterns obtained from ipsilateral or contralateral stimulation (Bl→Il, Bl→Cl). Likewise, we trained on ipsilateral or contralateral stimulation, and tested on bilateral stimulation (Il→Bl, Cl→Bl). As the time-resolved decoding analyses described above (**Time course of odor coding across stimulus laterality conditions**) suggested differential time courses of odor coding dependent on stimulus laterality, for the current set of analyses we trained and tested on data from all pairs of decoding time bins tiling the inhalation duration, resulting in a two-dimensional matrix of decoding results across training time (columns) and testing time (rows), which we refer to as the time-frequency similarity matrix (TFSM). Decoding accuracies along the diagonal of the matrix represent model performance when the model was trained and tested with data from the same decoding time bin. In contrast, values above the diagonal show the results when models trained on a prior time window were tested on future time windows, whereas values below the diagonal show results when models trained on a future time window were tested on past time windows.

To investigate the similarity of odor representations across stimulus laterality conditions, we took the difference of each pair of corresponding TFSMs, resulting in the following three TFSM contrasts: 1) Il→Cl − Cl→Il (Figure 5C), 2) Bl→Il − Il→Bl (Figure 6A), 3) Bl→Cl − Cl→Bl (Figure 6B). Prior to computing TFSM contrasts, the observed raw decoding accuracy values in each TFSM matrix (for Il, Cl, and Bl) were *z*-score normalized with respect to the corresponding null distributions generated for each electrode × training time bin × testing time bin as described above (**Decoding odor identity from odor-induced patterns of oscillatory activity**) by shuffling odor identity labels across trials (N = 500). We then computed TFSM contrast for each electrode. Statistical analysis of electrode-level TFSM contrasts mirrored the procedure described above (**Time course of odor coding across stimulus laterality conditions**), with the following differences: surrogate data to determine statistically significant temporal clusters was generated by shuffling data across either the training or testing time axes, one of which was selected randomly at each of the 500 iterations. Significance was assessed using a two-tailed threshold of greater than 0.5 s.d. or lesser than −0.5 s.d. at each training and testing time pair, and a two-tailed cluster-level threshold of *p* < 0.05.

### Distinguishability of odor representations between ipsilateral versus contralateral conditions

Representations of ipsilateral and contralateral odors were used to train and test a model to distinguish between the two stimulus laterality conditions (Figure 5D). To do this, we extracted time-frequency features from statistically significant coding epochs identified in the Il→Cl − Cl→Il contrast TFSM, during which a model trained on ipsilateral odors could classify contralateral odors, and vice versa. These temporal clusters spanned 18-27% of inhalation. To account for inter-subject variability in inhalation duration (note that the maximum inhalation duration across subjects was 2,662 ms), we chose to extract time-frequency features from a 600 ms window centered on the geometric centers of the two temporal clusters (50.5% for ipsilateral and 79% for contralateral). Classifier development and statistical analysis was performed as described above (**Decoding odor identity from odor-induced patterns of oscillatory activity**), with two differences: the null distribution was constructed by shuffling laterality labels, not odor type labels; laterality was classified between ipsilateral and contralateral odor trials matching trials of the same odor types.

### Time-resolved behavioral correlations of PC odor representations

We performed the behavioral correlations outlined above (**Decoding odor identity from odor-induced patterns of oscillatory activity)**, but in a time-resolved manner. This analysis was performed independently for each stimulus laterality condition, so as to determine whether the accuracy of PC odor representations would be correlated with subsequent behavioral performance on the odor identification task at different time points depending on stimulus laterality. Electrode-level decoding accuracies were again normalized with respect to null distributions constructed by shuffling labels.

For group-level results (Figure 7), at each resampling iteration, we computed each subject’s decoding accuracy by taking the average across the respective resampled electrodes, and then computed a Pearson’s correlation between decoding and behavioral accuracies across all subjects. Group-level results were obtained by taking the mean across 200 resampling procedures. To identify statistically significant temporal clusters of correlations, null distributions were constructed by shuffling the relative order of behavioral accuracies across subjects, and computing a Pearson’s correlations between the shuffled behavioral performance and decoding accuracy values (N = 500). Significance was again assessed using a one-tailed 1 s.d. pixel-level threshold and a one-tailed cluster-level threshold of *p* < 0.025 with respect to corresponding null distributions. For depiction of group-level results, the raw Pearson’s correlation values at each time point were *z*-score normalized with respect to null distributions constructed independently for each laterality condition. We note that the results were qualitatively similar when the Pearson’s correlation was computed after independently *z*-score normalizing both decoding and behavioral accuracies across subjects to achieve a normal distribution.

---

**Figure S1.**
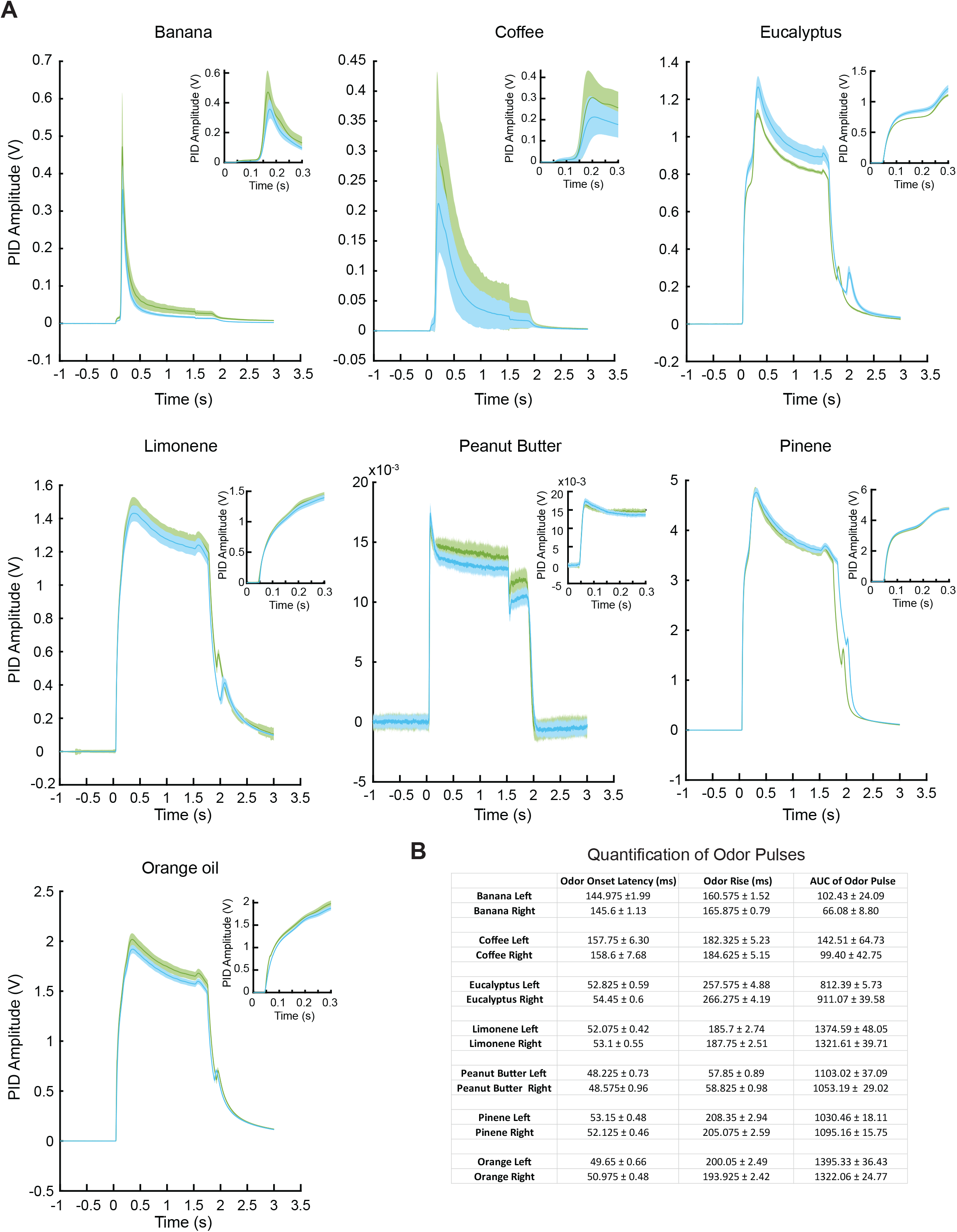
Odor pulse characteristics for left and right nostril delivery are matched. Related to Figure 1. **(A)** Photoionization detector (PID) characterization of each odor type delivered from the left and right valve sets. Each odor type was tested for 40 trials from the left and right delivery conditions following the setup described in Figure 1A. Two bottles of the same odorant were prepared (e.g. 20% pinene) and connected to the left and right nostril delivery setup (e.g. one bottle connected to valve 1 and the other connected to valve 7) ensuring that all valve combinations were tested across the seven unique odorants. Each panel shows the mean and standard deviation of all 40 trials from the left side (shown in green) overlaid with the mean and standard deviation of the 40 trials from the right (shown in blue) for a given odor type. The inset shows the zoomed in traces after valve opening (time = 0 s). For each odor type, the delivery of the odorant from the left versus the right valve setup resulted in nearly identical odor pulse characteristics. **(B)** Summary table of odor pulse characteristics shown in (A). Odor onset latency is defined as the time between valve opening and the odor pulse reaching 5% of its maximum amplitude. Odor rise time is defined as the time between valve opening and the odor pulse reaching 80% of its maximum amplitude. Area under the odor pulse is defined by the integral of the PID trace. The pulse characteristics of odorants delivered from the left versus right valves are matched. Note that natural odorants (banana, coffee, peanut butter) can have small variations in amplitude across the two bottles (i.e., left versus right) due to subtle differences in surface area of the odorant (i.e. 7 gram of banana in one bottle can have slightly more surface area than 7 gram of banana in the other bottle and result in slightly higher PID amplitude).

**Figure S2.**
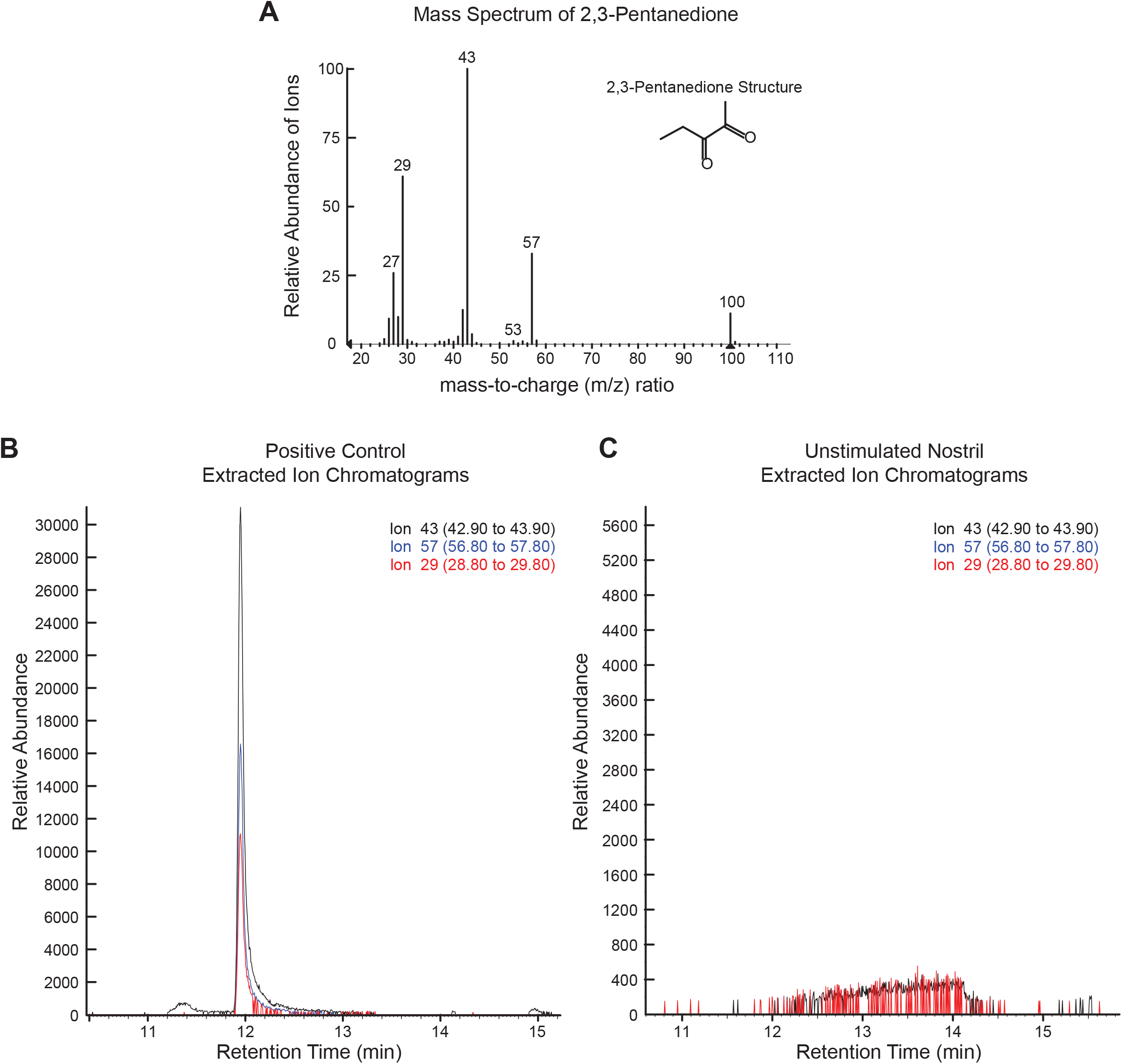
Odor delivery to one nostril does not contaminate the other nostril. Related to Figure 1. The odorant 2,3-pentanedione (0.05% concentration) was delivered to one nostril for 3 seconds while the unstimulated nostril was monitored for odor molecules using a SPME fiber. The odorant delivery followed the setup illustrated in Figure 1A. Prior to the odor delivery, the SPME fiber was inserted into the unstimulated nostril (the nostril that did not receive the odorant) and held in place until the end of odor delivery and the corresponding inhalation cycle. The SPME fiber was removed from the nostril at the end of inhalation and immediately tested for presence of 2,3-pentanedione using gas chromatography mass spectrometry (GC-MS). We selected 2,3-pentanedione due to its well-established characterization in olfactory experiments utilizing GC-MS. **(A)** Mass spectrum of 2,3-pentanedione according to the NIST Mass Spectral Library. The three most abundant fragment ions for odorant 2,3-pentanedione, m/z 43, m/z 29 and m/z 57, were used to examine the presence of the odor compound in (B) and (C). **(B)** Positive control. The SPME fiber was exposed to the 2,3-pentanedione (0.05% concentration) for 3 seconds to ensure that the exposure time was sufficient and that the odorant could be captured and detected with the GC-MS setup. As expected, all three ions were observed at a retention time of 12 minutes as shown by the alignment of the extracted ion chromatograms (EIC) of all three characteristic fragment ions. **(C)** Test protocol. As described above, the SPME fiber was held inside the unstimulated nostril while the other nostril received 2,3-pentanedione (0.05% concentration) for 3 seconds. The extracted ion chromatograms show that none of the three ions were observed at a retention time of 12 minutes, indicating that 2,3-pentanedione was not present at instrumentally detectable levels in the unstimulated nostril during inhalation.

**Figure S3.**
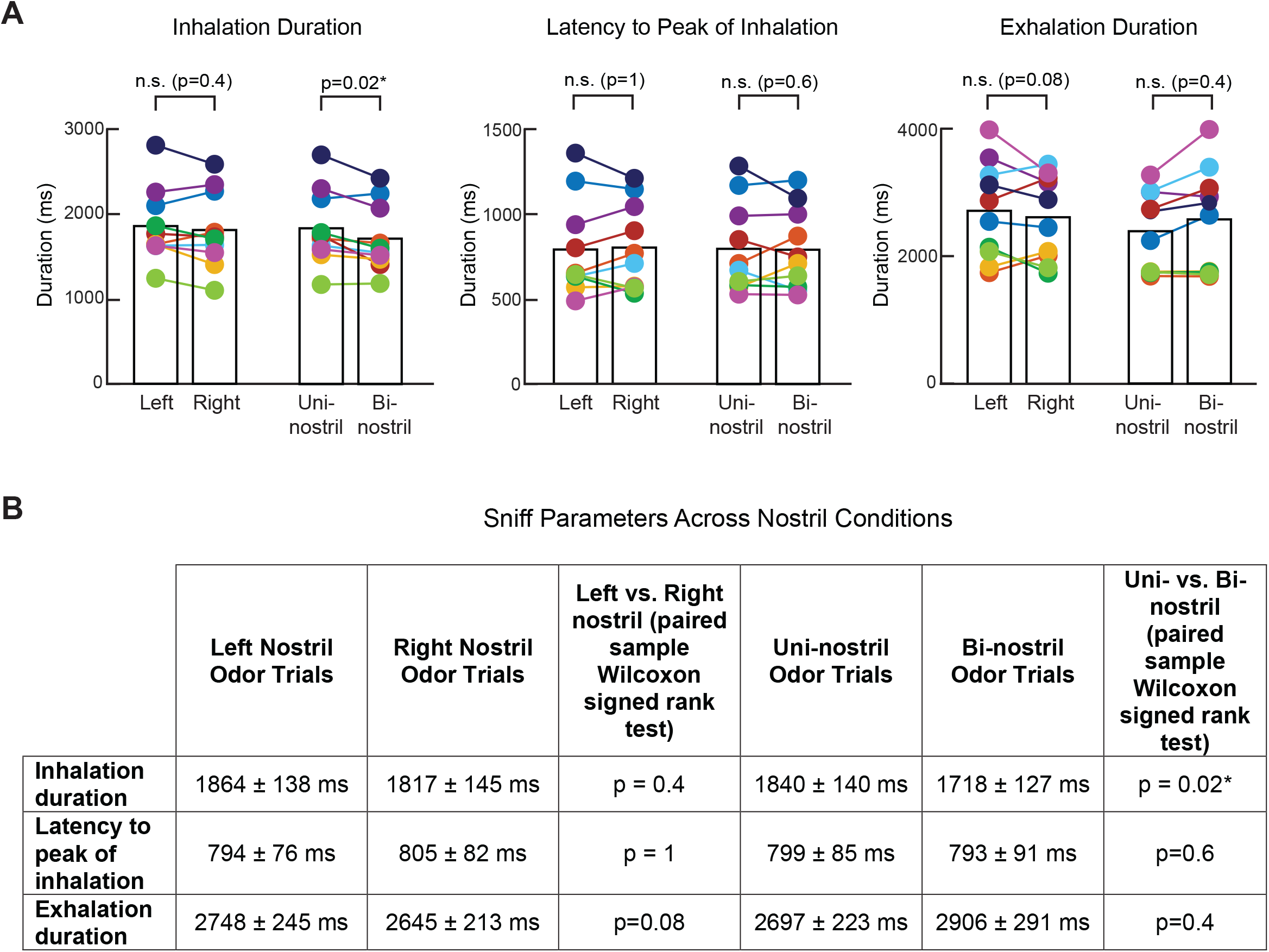
Sniff parameters across subjects and nostril conditions. Related to Figure 2. **(A)** Average inhalation duration, latency to peak of inhalation, and exhalation duration are shown for each subject and nostril condition. Each colored dot represents one subject. Left versus right nostril sniff parameters were not significantly different (paired sample Wilcoxon signed rank test). Inhalation duration was longer for uni-vs. bi-nostril odors but there were no significant differences between the two conditions for the other two parameters (paired sample Wilcoxon signed rank test, p-values shown on the graph). **(B)** Summary table lists subject-averaged sniff parameters for each nostril condition. Data shown as mean ± s.e.m.

**Figure S4.**
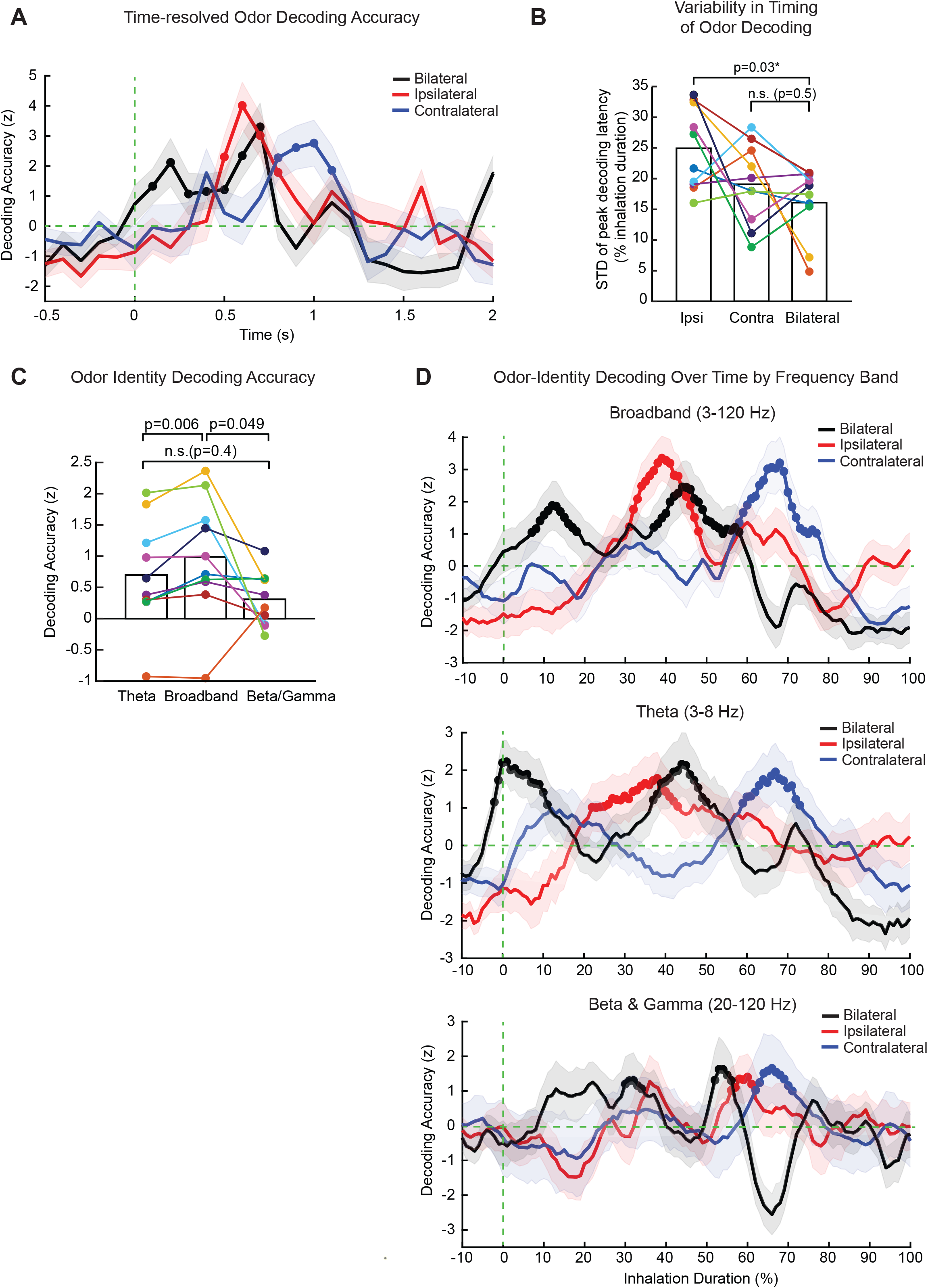
Ipsilateral and contralateral odors are encoded in temporally staggered single epochs while bi-nostril odors are encoded in two distinct temporal epochs. Related to Figure 4. **(A)** Subject-averaged time course of odor decoding for odors delivered via ipsilateral, contralateral, or bi-nostril routes depicted across raw time axis ([-0.5, 2] s with respect to sniff onset), as opposed to the time axis normalized with respect to subject-specific inhalation durations (as in Figure 4A). Solid lines correspond to the mean and the shaded lines represent the standard deviation of resampled distribution. Decoding accuracy values were first normalized with respect to odor label shuffled surrogate data, and then normalized with respect to time-shifted surrogate data. The latter null distribution was used to determine significant temporal clusters (one-tailed permutation test, cluster-level p<0.025), indicated with solid circles. **(B)** The temporal variability in the emergence of odor representations is not greater in the bi-nostril versus uni-nostril conditions. For each PC electrode, the latency to peak decoding accuracy of the first significant temporal cluster was computed separately for each of the three nostril conditions. Example electrode-level temporal patterns of odor coding are shown in Figure 4B. We then computed the standard deviation across all respective PC electrodes for each subject (colored dots), and compared these values at the group-level across nostril conditions. Ipsilateral condition had significantly greater temporal variability than the bilateral condition (paired-sample Wilcoxon signed rank test, p=0.03), whereas there was no difference between the contralateral and bilateral conditions (paired-sample Wilcoxon signed rank test, p=0.5). These data suggest that the dual coding observed in the bi-nostril condition in the group-level results (Figure 4A) cannot be explained by greater within-subject variability in the time course of odor coding across PC electrodes. **(C)** At the group-level, broadband oscillations had significantly better odor decoding accuracy compared to decoding with theta activity alone (paired-sample Wilcoxon signed rank test, p=0.006) or with beta/gamma activity alone (paired-sample Wilcoxon signed rank test, p=0.049). Each colored dot represents one subject. Classifiers were trained with data from the entire post-stimulus epoch using bi-nostril trials (as in Figures 3C, 3D). Electrode-level decoding accuracy values were first normalized with respect to odor label shuffled surrogate data, which were averaged across all respective electrode for each subject. **(D)** The time course of theta-based (middle panel) and beta/gamma-based (bottom panel) odor representations recapitulates the dynamics observed with broadband oscillations (shown in Figure 4A and also replicated in the top panel). Specifically, subject-averaged decoding accuracy time series for theta and beta-gamma similarly show that ipsilateral odors are encoded faster than contralateral odors, and that bi-nostril odor representations emerge in two distinct temporal clusters. Solid lines correspond to the mean and shaded lines represent the standard deviation of resampled distribution (see Methods). Decoding accuracy values were first normalized with respect to odor label shuffled surrogate data, and then normalized with respect to time-shifted surrogate data (see Methods). The latter null distribution was used to determine significant temporal clusters (one-tailed permutation test, cluster-level p<0.025), indicated with solid circles (see Methods). Time axis is shown as percentage of inhalation duration, where 0% and 100% represent the onset and end of inhalation, respectively.

**Figure S5.**
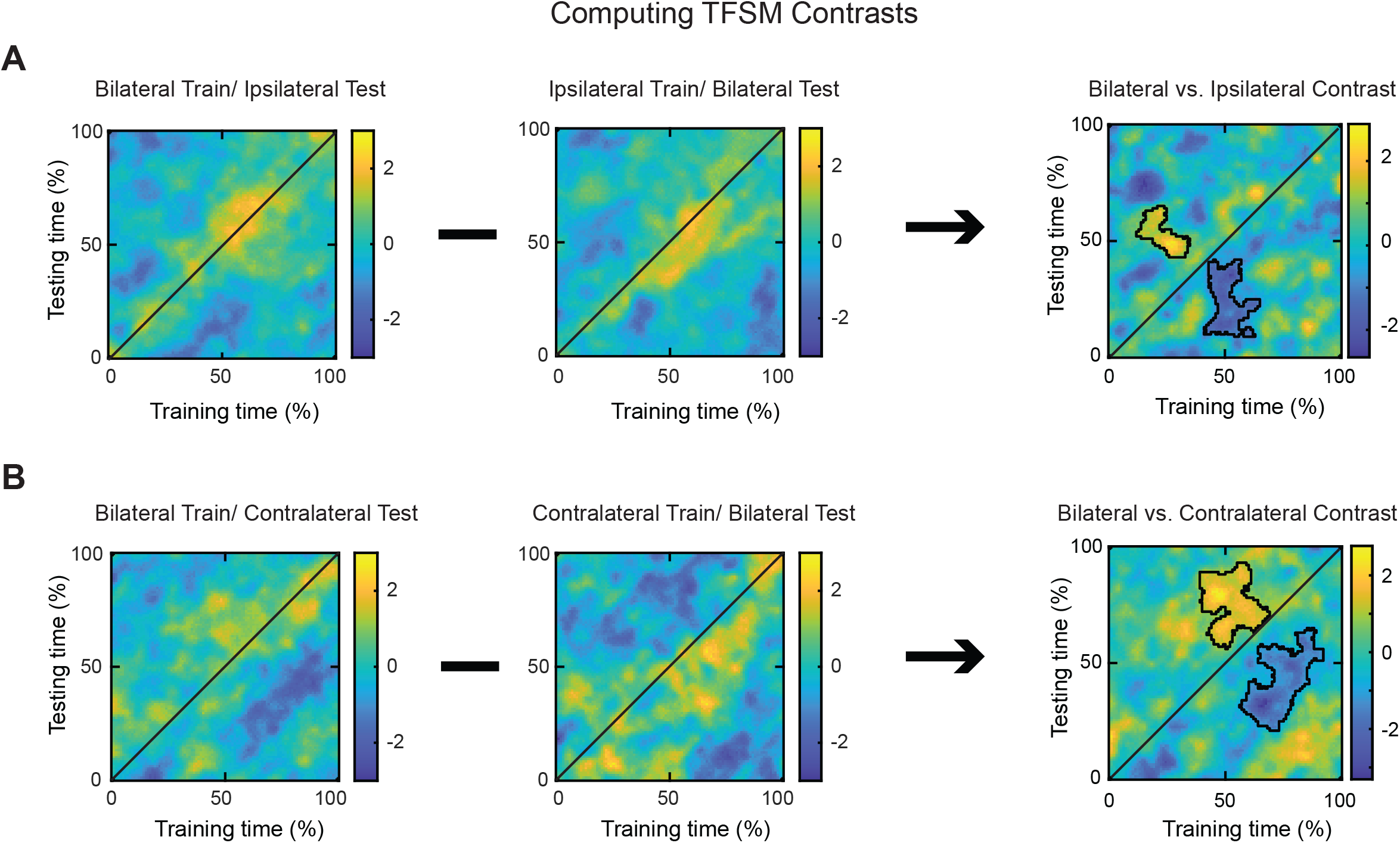
Illustration of time-frequency similarity analysis for bi-nostril vs. uni-nostril odor representations. Related to Figure 6. For each PC electrode, the time-frequency similarity matrices (TFSM) were computed by training the classifier on bilateral odor responses and testing on either ipsilateral **(A)** or contralateral **(B)** odor responses, across all possible training and testing time bins tiling the inhalation duration. The TFSM shown in (A) and (B) are averaged across all subjects. We also performed the same computation after reversing the training and testing sets (middle column). Decoding accuracy values were normalized with respect to odor label shuffled surrogate data. We then took the contrast of corresponding pairs of TFSMs (right column, also shown in Figure 6). Decoding accuracies in the contrast TFSM were further normalized with respect to surrogate data from random shifts across training and testing times, which was used to identify significant clusters (two-tailed permutation test, cluster-level p<0.05; see Methods), as marked with black contours. Time represents percentage of inhalation duration.

**Figure S6.**
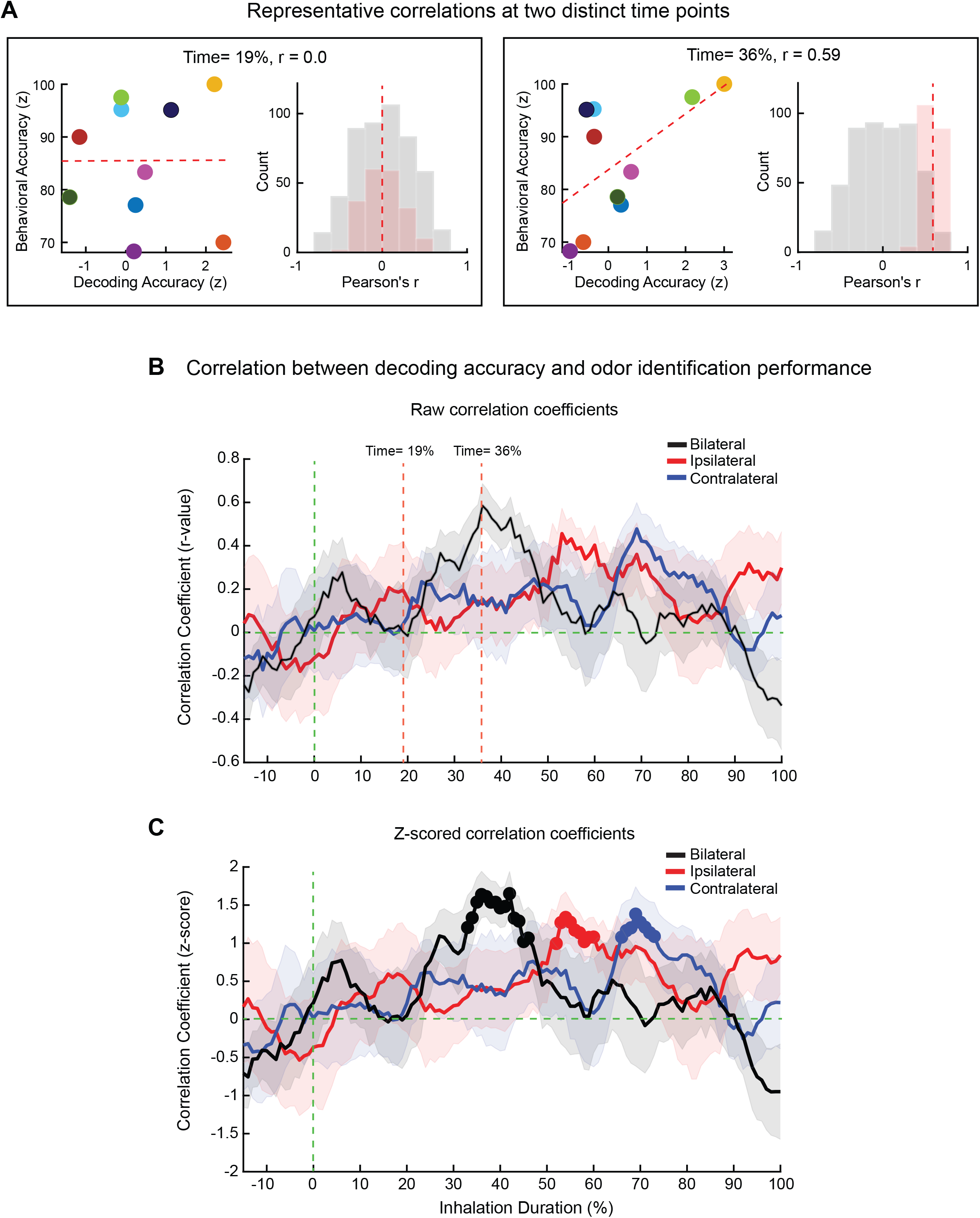
Methods schematic for computing time-resolved correlations between decoding accuracy and behavioral performance. Related to Figure 7. Subject-averaged time course of correlations between odor decoding accuracy and odor identification performance (shown in Figure 7, and also replicated in panel (C)). At each time point, Pearson’s correlation was computed between decoding accuracy and behavioral accuracy across subjects (as in Figure 3D). Data from two representative time points (19% and 36%) are shown for the bilateral condition in **(A)**. To control for variable number of PC electrodes across subjects, we used a resampling procedure. At each iteration, we randomly sampled the minimum number of PC electrodes across subjects, and computed each subject’s decoding accuracy by taking the average across the respective resampled electrodes. We then computed a Pearson’s correlation between decoding and behavioral accuracies across all subjects. The results from one such resampling iteration is shown in the left box in (A). The entire resampled distribution of correlation coefficients is shown in red (right box in (A)). The average correlation coefficient across the 200 resampling procedures is marked with dashed red lines, which was 0.0 at 19% of inhalation, and increased to 0.59 at 36%. Group-level results depicting this raw correlation coefficient across time for each of the three nostril conditions are shown in **(B)**, where the two representative time points are marked with orange dashed lines. Shaded areas represent standard deviation of the resampled distributions. To identify statistically significant temporal clusters of correlations, null distributions were constructed at each time point by shuffling the relative order of behavioral accuracies across subjects, and computing a Pearson’s correlations between the shuffled behavioral performance and decoding accuracy values (N = 500). These null distributions from the two representative time points are shown in grey in the right panels of (A). Significance was assessed using a one-tailed 1 s.d. pixel-level threshold and a one-tailed cluster-level threshold of p < 0.025 with respect to corresponding null distributions. Significant time clusters are indicated with solid circles in (C). Note that for the final depiction of group-level results in **(C)**, raw Pearson’s correlation values at each time point were z-score normalized with respect to the corresponding null distributions.

## REFERENCES

1. Rigolli, N., Magnoli, N., Rosasco, L., and Seminara, A. (2022). Learning to predict target location with turbulent odor plumes. eLife 11, e72196. 10.7554/eLife.72196.

2. Connor, E.G., McHugh, M.K., and Crimaldi, J.P. (2018). Quantification of airborne odor plumes using planar laser-induced fluorescence. Exp. Fluids 59, 137. 10.1007/s00348-018-2591-3.

3. Celani, A., Villermaux, E., and Vergassola, M. (2014). Odor Landscapes in Turbulent Environments. Phys. Rev. X 4, 041015. 10.1103/PhysRevX.4.041015.

4. Crimaldi, J.P., and Koseff, J.R. (2001). High-resolution measurements of the spatial and temporal scalar structure of a turbulent plume. Exp. Fluids 31, 90–102. 10.1007/s003480000263.

5. Porter, J., Craven, B., Khan, R.M., Chang, S.J., Kang, I., Judkewitz, B., Volpe, J., Settles, G., and Sobel, N. (2007). Mechanisms of scent-tracking in humans. Nat Neurosci 10, 27–29. 10.1038/nn1819.

6. Rajan, R., Clement, J.P., and Bhalla, U.S. (2006). Rats Smell in Stereo. Science 311, 666–670. 10.1126/science.1122096.

7. Gardiner, J.M., and Atema, J. (2010). The function of bilateral odor arrival time differences in olfactory orientation of sharks. Curr Biol 20, 1187–1191. 10.1016/j.cub.2010.04.053.

8. Catania, K.C. (2013). Stereo and serial sniffing guide navigation to an odour source in a mammal. Nat. Commun. 4, 1441. 10.1038/ncomms2444.

9. Wu, Y., Chen, K., Ye, Y., Zhang, T., and Zhou, W. (2020). Humans navigate with stereo olfaction. Proc. Natl. Acad. Sci. 117, 16065–16071. 10.1073/pnas.2004642117.

10. Gottfried, J.A. (2010). Central mechanisms of odour object perception. Nat. Rev. Neurosci. 11, 628– 641. 10.1038/nrn2883.

11. Uchida, N., Poo, C., and Haddad, R. (2013). Coding and Transformations in the Olfactory System. Annu. Rev. Neurosci. 37. 10.1146/annurev-neuro-071013-013941.

12. Brann, D.H., and Datta, S.R. (2020). Finding the Brain in the Nose. Annu. Rev. Neurosci. 43, 277–295. 10.1146/annurev-neuro-102119-103452.

13. Blazing, R.M., and Franks, K.M. (2020). Odor coding in piriform cortex: mechanistic insights into distributed coding. Curr. Opin. Neurobiol. 64, 96–102. 10.1016/j.conb.2020.03.001.

14. Isaacson, J.S. (2010). Odor representations in mammalian cortical circuits. Curr. Opin. Neurobiol. 20, 328–331. 10.1016/j.conb.2010.02.004.

15. Kay, L.M., Beshel, J., Brea, J., Martin, C., Rojas-Líbano, D., and Kopell, N. (2009). Olfactory oscillations: the what, how and what for. Trends Neurosci. 32, 207–214. 10.1016/j.tins.2008.11.008.

16. Dalal, T., Gupta, N., and Haddad, R. (2020). Bilateral and unilateral odor processing and odor perception. Commun Biol 3, 150. 10.1038/s42003-020-0876-6.

17. The synaptic organization of the brain, 5th ed (2004). (Oxford University Press) 10.1093/acprof:oso/9780195159561.001.1.

18. Kucharski, D., Burka, N., and Hall, W.G. (1990). The anterior limb of the anterior commissure is an access route to contralateral stored olfactory preference memories. Psychobiology 18, 195–204. 10.3758/BF03327227.

19. Russo, M.J., Franks, K.M., Oghaz, R., Axel, R., and Siegelbaum, S.A. (2020). Synaptic Organization of Anterior Olfactory Nucleus Inputs to Piriform Cortex. J Neurosci 40, 9414–9425. 10.1523/JNEUROSCI.0965-20.2020.

20. Grobman, M., Dalal, T., Lavian, H., Shmuel, R., Belelovsky, K., Xu, F., Korngreen, A., and Haddad, R. (2018). A Mirror-Symmetric Excitatory Link Coordinates Odor Maps across Olfactory Bulbs and Enables Odor Perceptual Unity. Neuron 99, 800–813 e6. 10.1016/j.neuron.2018.07.012.

21. Wilson, D.A. (1997). Binaral interactions in the rat piriform cortex. J Neurophysiol 78, 160–169. 10.1152/jn.1997.78.1.160.

22. Grimaud, J., Dorrell, W., Pehlevan, C., and Murthy, V. (2020). Bilateral Alignment of Receptive Fields in the Olfactory Cortex (Neuroscience) 10.1101/2020.02.24.960922.

23. Kucharski, D., and Hall, W.G. (1987). New Routes to Early Memories. Science 238, 786–788.

24. Savic, I., and Gulyas, B. (2000). PET shows that odors are processed both ipsilaterally and contralaterally to the stimulated nostril. NeuroReport 11, 2861–2866.

25. Porter, J., Anand, T., Johnson, B., Khan, R.M., and Sobel, N. (2005). Brain mechanisms for extracting spatial information from smell. Neuron 47, 581–592. 10.1016/j.neuron.2005.06.028.

26. Sobel, N., Khan, R.M., Hartley, C.A., Sullivan, E.V., and Gabrieli, J.D. (2000). Sniffing longer rather than stronger to maintain olfactory detection threshold. Chem Senses 25, 1–8.

27. Stettler, D.D., and Axel, R. (2009). Representations of Odor in the Piriform Cortex. Neuron 63, 854– 864. 10.1016/j.neuron.2009.09.005.

28. Roland, B., Deneux, T., Franks, K.M., Bathellier, B., and Fleischmann, A. (2017). Odor identity coding by distributed ensembles of neurons in the mouse olfactory cortex. eLife 6, e26337. 10.7554/eLife.26337.

29. Pashkovski, S.L., Iurilli, G., Brann, D., Chicharro, D., Drummey, K., Franks, K.M., Panzeri, S., and Datta, S.R. (2020). Structure and flexibility in cortical representations of odour space. Nature 583, 253–258. 10.1038/s41586-020-2451-1.

30. Schoonover, C.E., Ohashi, S.N., Axel, R., and Fink, A.J.P. (2021). Representational drift in primary olfactory cortex. Nature 594, 541–546. 10.1038/s41586-021-03628-7.

31. Jiang, H., Schuele, S., Rosenow, J., Zelano, C., Parvizi, J., Tao, J.X., Wu, S., and Gottfried, J.A. (2017). Theta Oscillations Rapidly Convey Odor-Specific Content in Human Piriform Cortex. Neuron 94, 207–219.e4. 10.1016/j.neuron.2017.03.021.

32. Yang, A.I., Dikecligil, G.N., Jiang, H., Das, S.R., Stein, J.M., Schuele, S.U., Rosenow, J.M., Davis, K.A., Lucas, T.H., and Gottfried, J.A. (2021). The what and when of olfactory working memory in humans. Curr. Biol. 31, 4499–4511.e8. 10.1016/j.cub.2021.08.004.

33. Yang, Q., Zhou, G., Noto, T., Templer, J.W., Schuele, S.U., Rosenow, J.M., Lane, G., and Zelano, C. (2022). Smell-induced gamma oscillations in human olfactory cortex are required for accurate perception of odor identity. PLOS Biol. 20, e3001509. 10.1371/journal.pbio.3001509.

34. Zhou, G., Lane, G., Noto, T., Arabkheradmand, G., Gottfried, J.A., Schuele, S.U., Rosenow, J.M., Olofsson, J.K., Wilson, D.A., and Zelano, C. (2019). Human olfactory-auditory integration requires phase synchrony between sensory cortices. Nat. Commun. 10, 1168. 10.1038/s41467-019-09091-3.

35. Small, D.M., Gerber, J.C., Mak, Y.E., and Hummel, T. (2005). Differential Neural Responses Evoked by Orthonasal versus Retronasal Odorant Perception in Humans. Neuron 47, 593–605. 10.1016/j.neuron.2005.07.022.

36. Kahana-Zweig, R., Geva-Sagiv, M., Weissbrod, A., Secundo, L., Soroker, N., and Sobel, N. (2016). Measuring and Characterizing the Human Nasal Cycle. PLoS One 11, e0162918. 10.1371/journal.pone.0162918.

37. Hasegawa, M., and Kern, E.B. (1977). The human nasal cycle. Mayo Clin. Proc. 52, 28–34.

38. Tonoike, M., Yamaguchi, M., Kaetsu, I., Kida, H., Seo, R., and Koizuka, I. (1998). Ipsilateral dominance of human olfactory activated centers estimated from event-related magnetic fields measured by 122-channel whole-head neuromagnetometer using odorant stimuli synchronized with respirations. Ann N Acad Sci 855, 579–590.

39. Lascano, A.M., Hummel, T., Lacroix, J.S., Landis, B.N., and Michel, C.M. (2010). Spatio-temporal dynamics of olfactory processing in the human brain: an event-related source imaging study. Neuroscience 167, 700–708. 10.1016/j.neuroscience.2010.02.013.

40. Stadlbauer, A., Kaltenhauser, M., Buchfelder, M., Brandner, S., Neuhuber, W.L., and Renner, B. (2016). Spatiotemporal Pattern of Human Cortical and Subcortical Activity during Early-Stage Odor Processing. Chem Senses 41, 783–794. 10.1093/chemse/bjw074.

41. Grothe, B., Pecka, M., and McAlpine, D. (2010). Mechanisms of Sound Localization in Mammals. Physiol. Rev. 90, 983–1012. 10.1152/physrev.00026.2009.

42. Smith, S.M., Jenkinson, M., Woolrich, M.W., Beckmann, C.F., Behrens, T.E.J., Johansen-Berg, H., Bannister, P.R., De Luca, M., Drobnjak, I., Flitney, D.E., et al. (2004). Advances in functional and structural MR image analysis and implementation as FSL. NeuroImage 23, S208–S219. 10.1016/j.neuroimage.2004.07.051.

43. Brainard, D.H. (1997). The Psychophysics Toolbox. Spat. Vis. 10, 433–436. 10.1163/156856897X00357.

44. Delorme, A., and Makeig, S. (2004). EEGLAB: an open source toolbox for analysis of single-trial EEG dynamics including independent component analysis. J. Neurosci. Methods 134, 9–21. 10.1016/j.jneumeth.2003.10.009.

45. Oostenveld, R., Fries, P., Maris, E., and Schoffelen, J.-M. (2011). FieldTrip: open source software for advanced analysis of MEG, EEG, and invasive electrophysiological data. Comput. Intell. Neurosci. 2011, 1:1–1:9. 10.1155/2011/156869.

46. Chang, C.-C., and Lin, C.-J. (2011). LIBSVM: A library for support vector machines. ACM Trans. Intell. Syst. Technol. 2, 27:1–27:27. 10.1145/1961189.1961199.

47. Li, G., Jiang, S., Paraskevopoulou, S.E., Wang, M., Xu, Y., Wu, Z., Chen, L., Zhang, D., and Schalk, G. (2018). Optimal referencing for stereo-electroencephalographic (SEEG) recordings. NeuroImage 183, 327–335. 10.1016/j.neuroimage.2018.08.020.

48. Staresina, B.P., Bergmann, T.O., Bonnefond, M., van der Meij, R., Jensen, O., Deuker, L., Elger, C.E., Axmacher, N., and Fell, J. (2015). Hierarchical nesting of slow oscillations, spindles and ripples in the human hippocampus during sleep. Nat. Neurosci. 18, 1679–1686. 10.1038/nn.4119.

49. Maris, E., and Oostenveld, R. (2007). Nonparametric statistical testing of EEG- and MEG-data. J. Neurosci. Methods 164, 177–190. 10.1016/j.jneumeth.2007.03.024.

