## Supplemental Figures for "Odor representations from the two nostrils are temporally segregated in human piriform cortex"

A

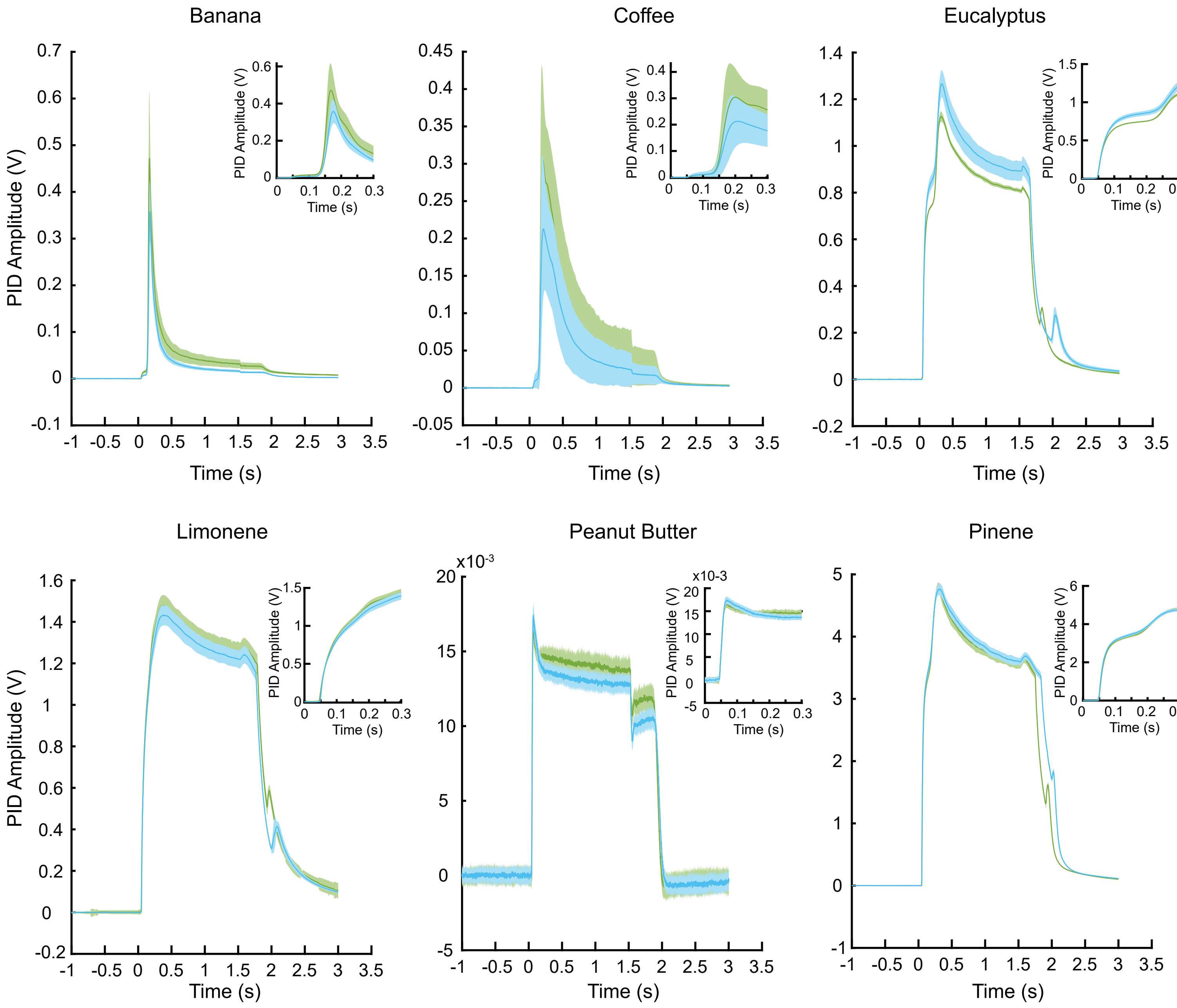

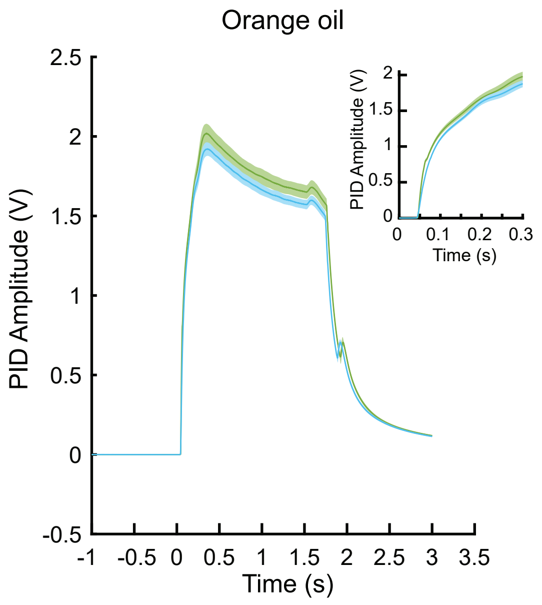

B

Quantification of Odor Pulses

| | Odor Onset Latency (ms) | Odor Rise (ms) | AUC of Odor Pulse |
|---|---|---|---|
| **Banana Left** | 144.975 ±1.99 | 160.575 ± 1.52 | 102.43 ± 24.09 |
| **Banana Right** | 145.6 ± 1.13 | 165.875 ± 0.79 | 66.08 ± 8.80 |
| **Coffee Left** | 157.75 ± 6.30 | 182.325 ± 5.23 | 142.51 ± 64.73 |
| **Coffee Right** | 158.6 ± 7.68 | 184.625 ± 5.15 | 99.40 ± 42.75 |
| **Eucalyptus Left** | 52.825 ± 0.59 | 257.575 ± 4.88 | 812.39 ± 5.73 |
| **Eucalyptus Right** | 54.45 ± 0.6 | 266.275 ± 4.19 | 911.07 ± 39.58 |
| **Limonene Left** | 52.075 ± 0.42 | 185.7 ± 2.74 | 1374.59 ± 48.05 |
| **Limonene Right** | 53.1 ± 0.55 | 187.75 ± 2.51 | 1321.61 ± 39.71 |
| **Peanut Butter Left** | 48.225 ± 0.73 | 57.85 ± 0.89 | 1103.02 ± 37.09 |
| **Peanut Butter Right** | 48.575± 0.96 | 58.825 ± 0.98 | 1053.19 ± 29.02 |
| **Pinene Left** | 53.15 ± 0.48 | 208.35 ± 2.94 | 1030.46 ± 18.11 |
| **Pinene Right** | 52.125 ± 0.46 | 205.075 ± 2.59 | 1095.16 ± 15.75 |
| **Orange Left** | 49.65 ± 0.66 | 200.05 ± 2.49 | 1395.33 ± 36.43 |
| **Orange Right** | 50.975 ± 0.48 | 193.925 ± 2.42 | 1322.06 ± 24.77 |

**Figure S1. Odor pulse characteristics for left and right nostril delivery are matched. Related to Figure 1.**

**(A)** Photoionization detector (PID) characterization of each odor type delivered from the left and right valve sets. Each odor type was tested for 40 trials from the left and right delivery conditions following the setup described in Figure 1A. Two bottles of the same odorant were prepared (e.g. 20% pinene) and connected to the left and right nostril delivery setup (e.g. one bottle connected to valve 1 and the other connected to valve 7) ensuring that all valve combinations were tested across the seven unique odorants. Each panel shows the mean and standard deviation of all 40 trials from the left side (shown in green) overlaid with the mean and standard deviation of the 40 trials from the right (shown in blue) for a given odor type. The inset shows the zoomed in traces after valve opening (time = 0 s). For each odor type, the delivery of the odorant from the left versus the right valve setup resulted in nearly identical odor pulse characteristics. **(B)** Summary table of odor pulse characteristics shown in (A). Odor onset latency is defined as the time between valve opening and the odor pulse reaching 5% of its maximum amplitude. Odor rise time is defined as the time between valve opening and the odor pulse reaching 80% of its maximum amplitude. Area under the odor pulse is defined by the integral of the PID trace. The pulse characteristics of odorants delivered from the left versus right valves are matched. Note that natural odorants (banana, coffee, peanut butter) can have small variations in amplitude across the two bottles (i.e., left versus right) due to subtle differences in surface area of the odorant (i.e. 7 gram of banana in one bottle can have slightly more surface area than 7 gram of banana in the other bottle and result in slightly higher PID amplitude).

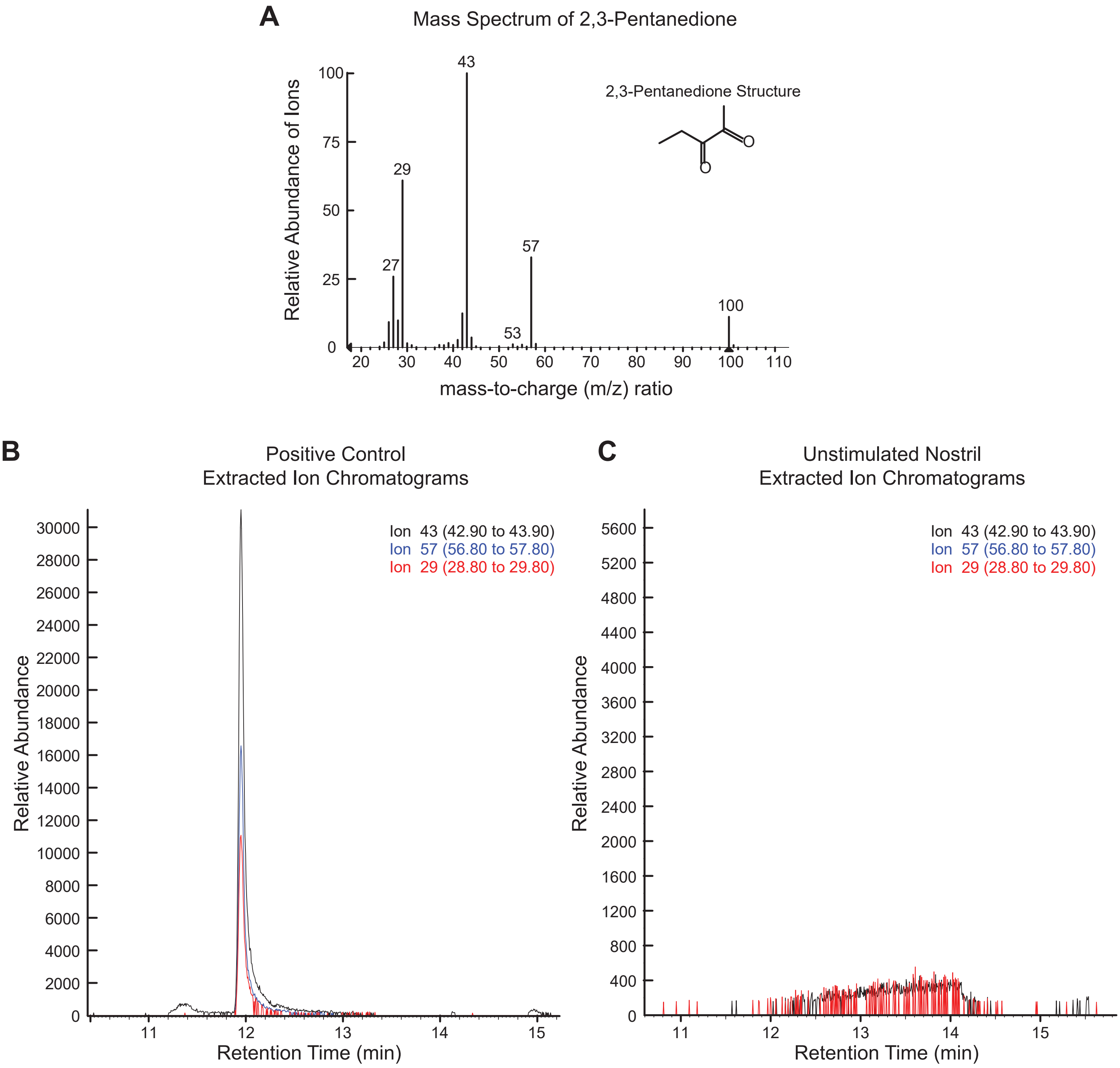

**Figure S2. Odor delivery to one nostril does not contaminate the other nostril. Related to Figure 1.**

The odorant 2,3-pentanedione (0.05% concentration) was delivered to one nostril for 3 seconds while the unstimulated nostril was monitored for odor molecules using a SPME fiber. The odorant delivery followed the setup illustrated in Figure 1A. Prior to the odor delivery, the SPME fiber was inserted into the unstimulated nostril (the nostril that did not receive the odorant) and held in place until the end of odor delivery and the corresponding inhalation cycle. The SPME fiber was removed from the nostril at the end of inhalation and immediately tested for presence of 2,3-pentanedione using gas chromatography mass spectrometry (GC-MS). We selected 2,3-pentanedione due to its well-established characterization in olfactory experiments utilizing GC-MS. **(A)** Mass spectrum of 2,3-pentanedione according to the NIST Mass Spectral Library. The three most abundant fragment ions for odorant 2,3-pentanedione, m/z 43, m/z 29 and m/z 57, were used to examine the presence of the odor compound in (B) and (C). **(B)** Positive control. The SPME fiber was exposed to the 2,3-pentanedione (0.05% concentration) for 3 seconds to ensure that the exposure time was sufficient and that the odorant could be captured and detected with the GC-MS setup. As expected, all three ions were observed at a retention time of 12 minutes as shown by the alignment of the extracted ion chromatograms (EIC) of all three characteristic fragment ions. **(C)** Test protocol. As described above, the SPME fiber was held inside the unstimulated nostril while the other nostril received 2,3-pentanedione (0.05% concentration) for 3 seconds. The extracted ion chromatograms show that none of the three ions were observed at a retention time of 12 minutes, indicating that 2,3-pentanedione was not present at instrumentally detectable levels in the unstimulated nostril during inhalation.

**A**

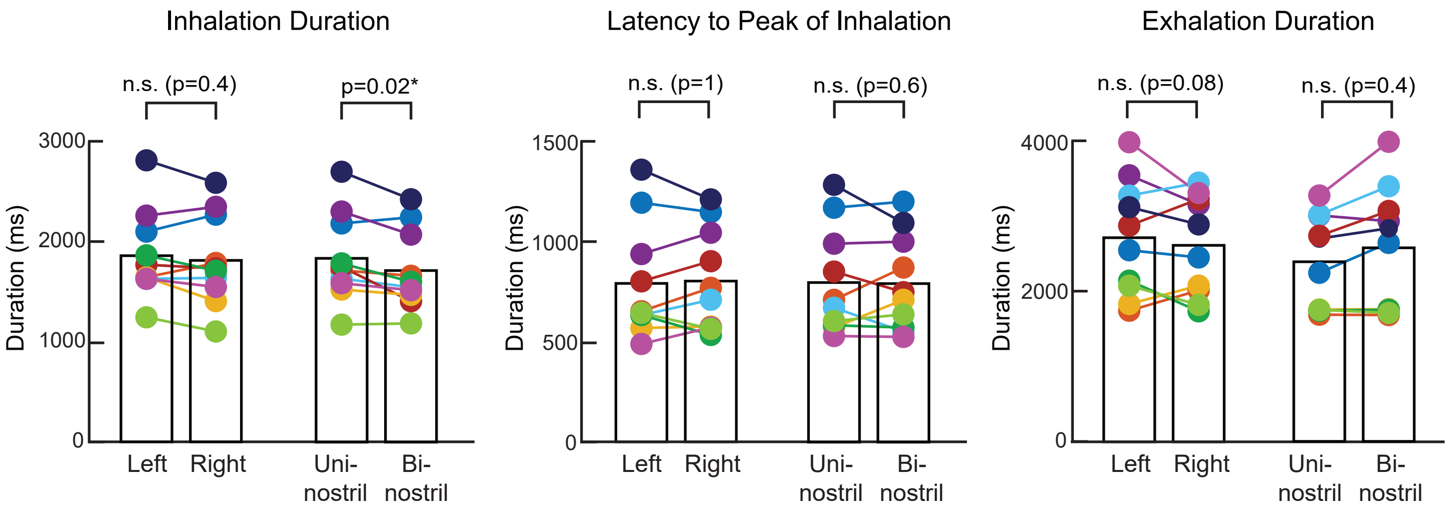

**B**

Sniff Parameters Across Nostril Conditions

| | Left Nostril Odor Trials | Right Nostril Odor Trials | Left vs. Right nostril (paired sample Wilcoxon signed rank test) | Uni-nostril Odor Trials | Bi-nostril Odor Trials | Uni- vs. Bi-nostril (paired sample Wilcoxon signed rank test) |
|---|---|---|---|---|---|---|
| **Inhalation duration** | 1864 ± 138 ms | 1817 ± 145 ms | p = 0.4 | 1840 ± 140 ms | 1718 ± 127 ms | p = 0.02* |
| **Latency to peak of inhalation** | 794 ± 76 ms | 805 ± 82 ms | p = 1 | 799 ± 85 ms | 793 ± 91 ms | p=0.6 |
| **Exhalation duration** | 2748 ± 245 ms | 2645 ± 213 ms | p=0.08 | 2697 ± 223 ms | 2906 ± 291 ms | p=0.4 |

**Figure S3. Sniff parameters across subjects and nostril conditions. Related to Figure 2.**

**(A)** Average inhalation duration, latency to peak of inhalation, and exhalation duration are shown for each subject and nostril condition. Each colored dot represents one subject. Left versus right nostril sniff parameters were not significantly different (paired sample Wilcoxon signed rank test). Inhalation duration was longer for uni- vs. bi-nostril odors but there were no significant differences between the two conditions for the other two parameters (paired sample Wilcoxon signed rank test, p-values shown on the graph). **(B)** Summary table lists subject-averaged sniff parameters for each nostril condition. Data shown as mean ± s.e.m.

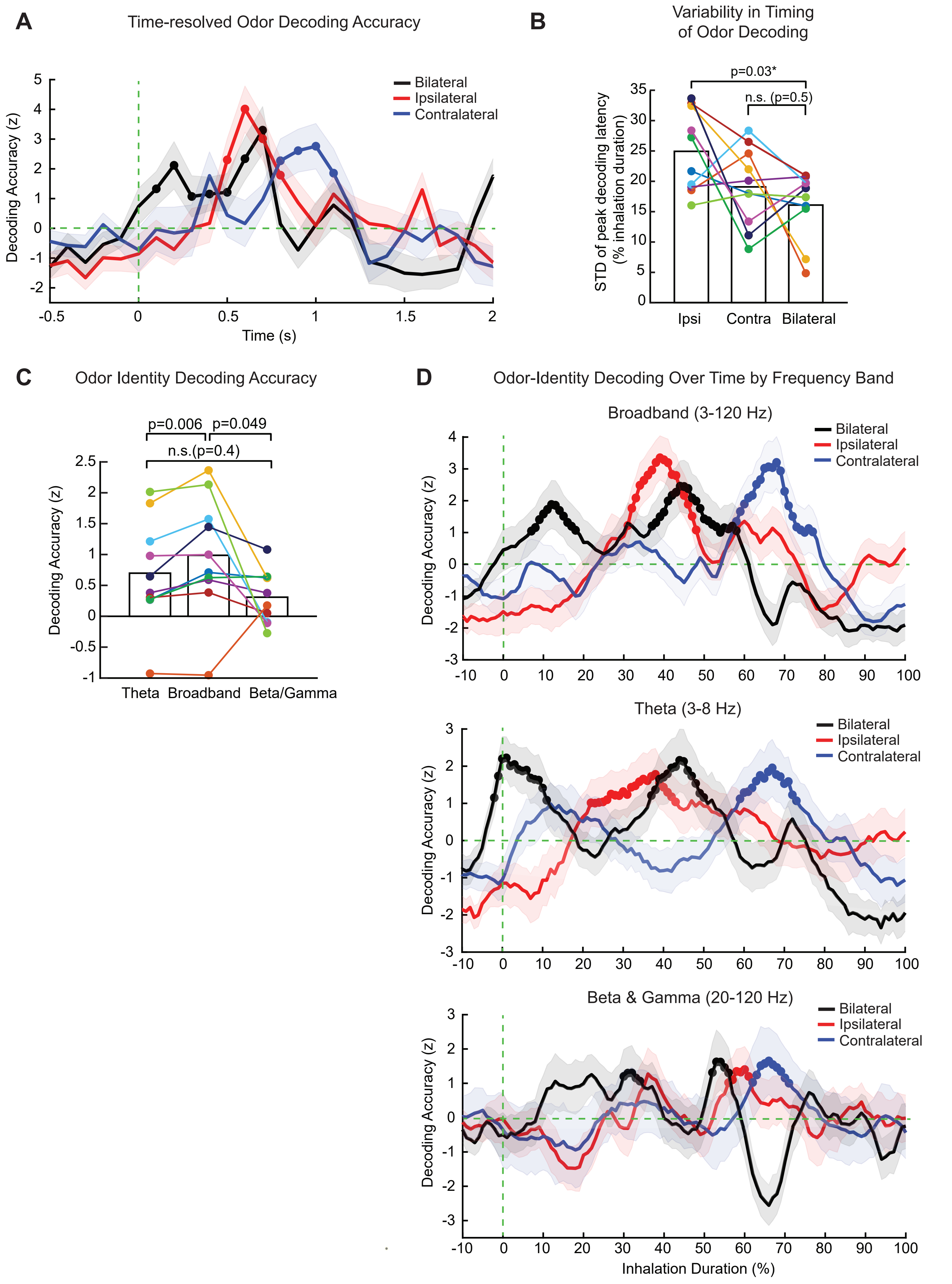
A
Time-resolved Odor Decoding Accuracy
Bilateral
Ipsilateral
Contralateral
Decoding Accuracy (z)
Time (s)
B
Variability in Timing of Odor Decoding
p=0.03*
n.s. (p=0.5)
STD of peak decoding latency (% inhalation duration)
Ipsi
Contra
Bilateral
C
Odor Identity Decoding Accuracy
p=0.006
p=0.049
n.s.(p=0.4)
Decoding Accuracy (z)
Theta
Broadband
Beta/Gamma
D
Odor-Identity Decoding Over Time by Frequency Band
Broadband (3-120 Hz)
Theta (3-8 Hz)
Beta & Gamma (20-120 Hz)
Inhalation Duration (%)

**Figure S4. Ipsilateral and contralateral odors are encoded in temporally staggered single epochs while bi-nostril odors are encoded in two distinct temporal epochs. Related to Figure 4.**

**(A)** Subject-averaged time course of odor decoding for odors delivered via ipsilateral, contralateral, or bi-nostril routes depicted across raw time axis ([-0.5, 2] s with respect to sniff onset), as opposed to the time axis normalized with respect to subject-specific inhalation durations (as in Figure 4A). Solid lines correspond to the mean and the shaded lines represent the standard deviation of resampled distribution. Decoding accuracy values were first normalized with respect to odor label shuffled surrogate data, and then normalized with respect to time-shifted surrogate data. The latter null distribution was used to determine significant temporal clusters (one-tailed permutation test, cluster-level $p<0.025$), indicated with solid circles. **(B)** The temporal variability in the emergence of odor representations is not greater in the bi-nostril versus uni-nostril conditions. For each PC electrode, the latency to peak decoding accuracy of the first significant temporal cluster was computed separately for each of the three nostril conditions. Example electrode-level temporal patterns of odor coding are shown in Figure 4B. We then computed the standard deviation across all respective PC electrodes for each subject (colored dots), and compared these values at the group-level across nostril conditions. Ipsilateral condition had significantly greater temporal variability than the bilateral condition (paired-sample Wilcoxon signed rank test, $p=0.03$), whereas there was no difference between the contralateral and bilateral conditions (paired-sample Wilcoxon signed rank test, $p=0.5$). These data suggest that the dual coding observed in the bi-nostril condition in the group-level results (Figure 4A) cannot be explained by greater within-subject variability in the time course of odor coding across PC electrodes. **(C)** At the group-level, broadband oscillations had significantly better odor decoding accuracy compared to decoding with theta activity alone (paired-sample Wilcoxon signed rank test, $p=0.006$) or with beta/gamma activity alone (paired-sample Wilcoxon signed rank test, $p=0.049$). Each colored dot represents one subject. Classifiers were trained with data from the entire post-stimulus epoch using bi-nostril trials (as in Figures 3C, 3D). Electrode-level decoding accuracy values were first normalized with respect to odor label shuffled surrogate data, which were averaged across all respective electrode for each subject. **(D)** The time course of theta-based (middle panel) and beta/gamma-based (bottom panel) odor representations recapitulates the dynamics observed with broadband oscillations (shown in Figure 4A and also replicated in the top panel). Specifically, subject-averaged decoding accuracy time series for theta and beta-gamma similarly show that ipsilateral odors are encoded faster than contralateral odors, and that bi-nostril odor representations emerge in two distinct temporal clusters. Solid lines correspond to the mean and shaded lines represent the standard deviation of resampled distribution (see Methods). Decoding accuracy values were first normalized with respect to odor label shuffled surrogate data, and then normalized with respect to time-shifted surrogate data (see Methods). The latter null distribution was used to determine significant temporal clusters (one-tailed permutation test, cluster-level $p<0.025$), indicated with solid circles (see Methods). Time axis is shown as percentage of inhalation duration, where 0% and 100% represent the onset and end of inhalation, respectively.

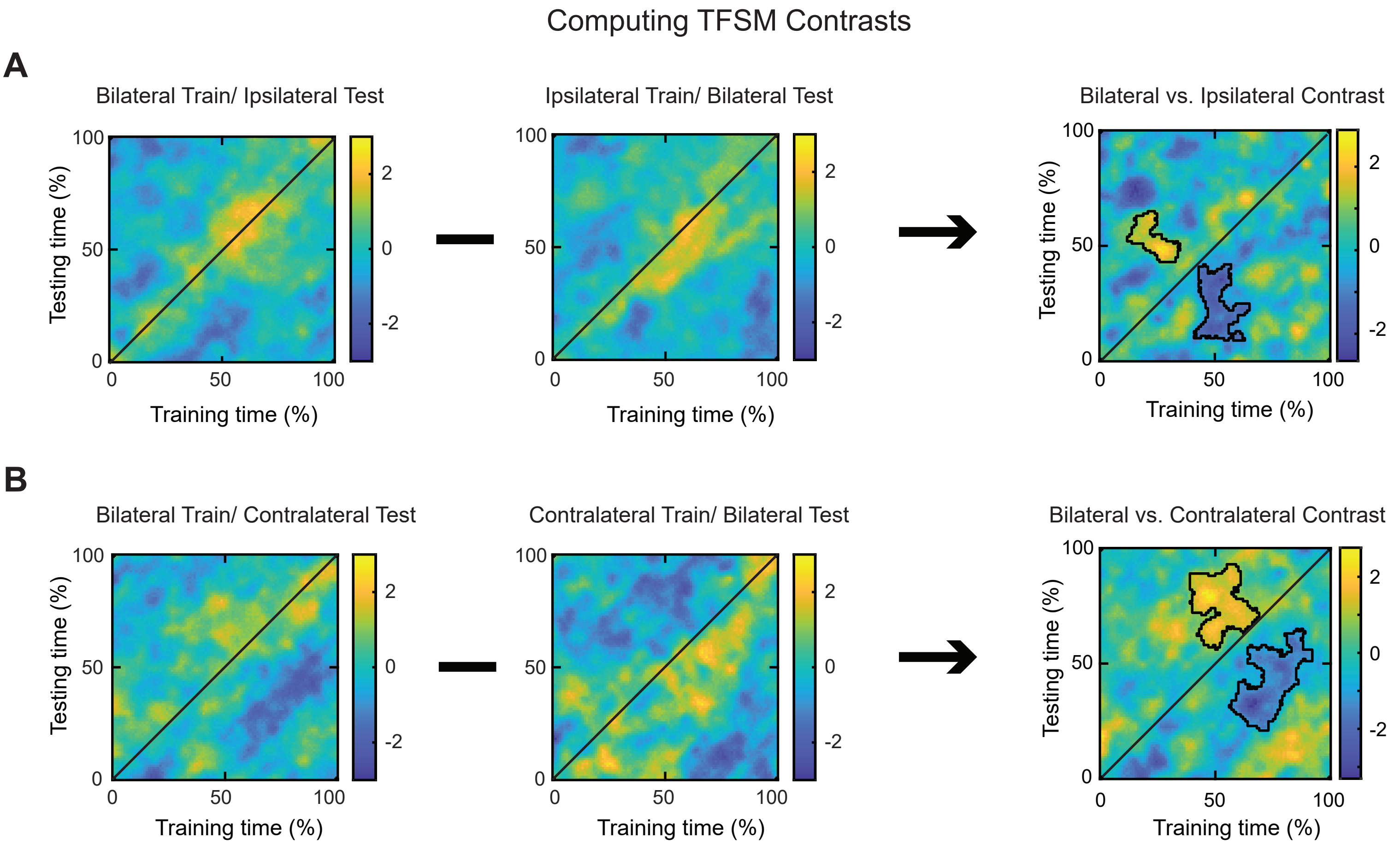

**Figure S5. Illustration of time-frequency similarity analysis for bi-nostril vs. uni-nostril odor representations. Related to Figure 6.**

For each PC electrode, the time-frequency similarity matrices (TFSM) were computed by training the classifier on bilateral odor responses and testing on either ipsilateral **(A)** or contralateral **(B)** odor responses, across all possible training and testing time bins tiling the inhalation duration. The TFSM shown in (A) and (B) are averaged across all subjects. We also performed the same computation after reversing the training and testing sets (middle column). Decoding accuracy values were normalized with respect to odor label shuffled surrogate data. We then took the contrast of corresponding pairs of TFSMs (right column, also shown in Figure 6). Decoding accuracies in the contrast TFSM were further normalized with respect to surrogate data from random shifts across training and testing times, which was used to identify significant clusters (two-tailed permutation test, cluster-level $p<0.05$; see Methods), as marked with black contours. Time represents percentage of inhalation duration.

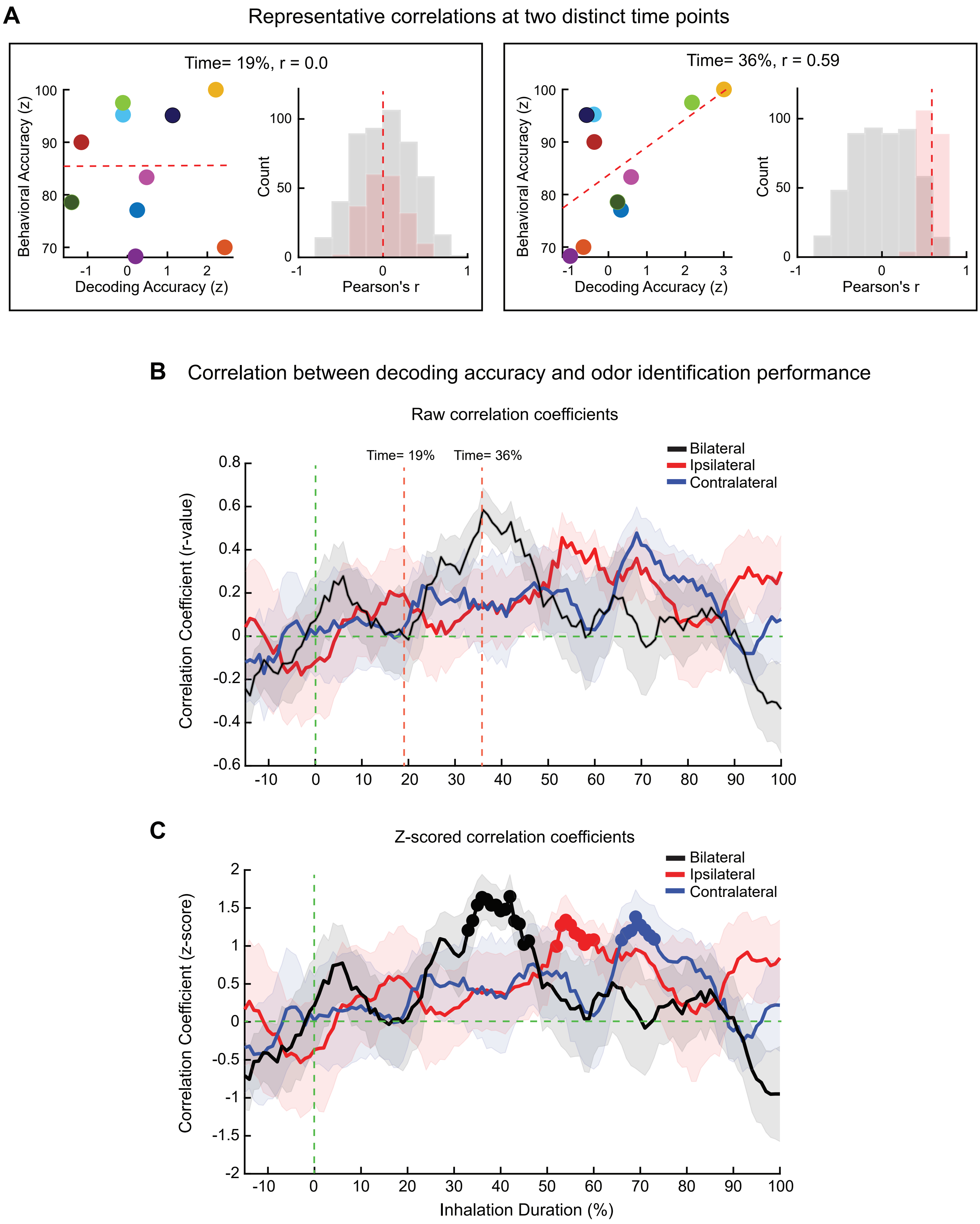

A
Representative correlations at two distinct time points
Time= 19%, r = 0.0
Behavioral Accuracy (z)
Decoding Accuracy (z)
Count
Pearson's r
Time= 36%, r = 0.59
B
Correlation between decoding accuracy and odor identification performance
Raw correlation coefficients
Time= 19%
Time= 36%
Bilateral
Ipsilateral
Contralateral
Correlation Coefficient (r-value)
C
Z-scored correlation coefficients
Correlation Coefficient (z-score)
Inhalation Duration (%)
